# Condensin I subunit Cap-G is essential for proper gene expression during the maturation of post-mitotic neurons

**DOI:** 10.1101/2020.01.14.904409

**Authors:** Amira Hassan, Pablo Araguas Rodriguez, Stefan K. Heidmann, Emma L. Walmsley, Gabriel N. Aughey, Tony D. Southall

**Affiliations:** Department of Life Sciences, Imperial College London, Sir Ernst Chain Building, London, UK; Lehrstuhl für Genetik, University of Bayreuth, Bayreuth, Germany

## Abstract

The condensin complex is essential for mitotic chromosome assembly and segregation during cell divisions, however, little is known about its function in post-mitotic, differentiated cells. Here we report a novel role for the condensin I subunit Cap-G in *Drosophila* neurons. We show that, despite not requiring condensin for mitotic chromosome compaction, post-mitotic neurons express Cap-G and that knockdown of Cap-G specifically in neurons (from their birth onwards) results in developmental arrest, behavioural defects, and dramatic gene expression changes. These include reduced expression of a subset of neuronal genes and aberrant expression of genes that are not normally expressed in the developing brain. Knockdown of Cap-G in more mature neurons also results in similar phenotypes but to a lesser degree. Furthermore, we see dynamic binding of Cap-G to chromatin at distinct loci in neural stem cells and differentiated neurons. Therefore, Cap-G is essential for proper gene expression in neurons and plays an important role during the early stages of neuronal development.

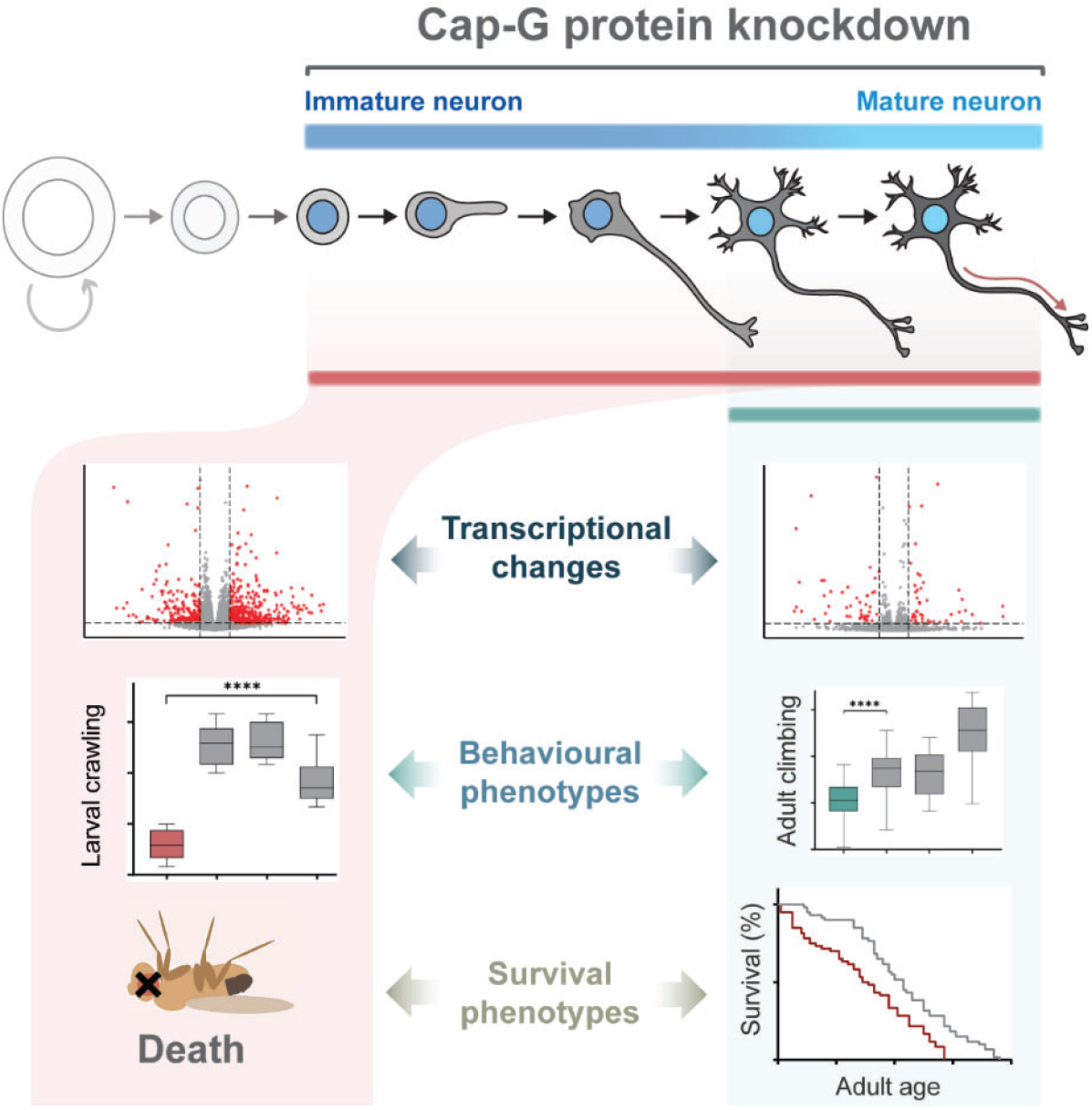

## Introduction

Differentiated neurons are post-mitotic cells – they lack the ability to further divide to produce daughter cells. However, newly born neurons are not immediately ready to synapse with other neurons, nor generate action potentials. These immature cells undergo morphological changes, generating dendrites and axons, which will eventually form synapses with target cell(s) [1]. Furthermore, newly born neurons display a level of developmental plasticity that is not apparent in terminally differentiated neurons, having been shown to have the potential to dedifferentiate under conditions that do not affect more mature cells [2, 3]. However, underlying changes at the chromatin level are only just starting to be investigated. Changes in gene expression, chromatin accessibility and the 3D genome organisation all occur as neurons differentiate from progenitor cells [4–9]. Currently, little is known about the mechanisms underlying this transition and the molecular factors that coordinate it.

During mitosis eukaryotic DNA is packaged into condensed chromosomes for efficient segregation between daughter cells. The formation of these highly organised chromosome structures is achieved largely by the actions of the condensin complex [10]. Components of the condensin complex form a loop through which DNA is constrained under the regulation of ATP-dependent SMC subunits in a similar manner to the related cohesin complex [10]. Most eukaryotes are thought to have two condensin complexes - condensin I and II. Both condensin complexes share a heterodimer of SMC subunits (SMC2/4) and differ based on inclusion of paralogous kleisin subunits (Cap-H/H2), and two HEAT-repeat subunits (CAP-G/G2 and CAP-D2/D3) [11] (Figure 1A). Each condensin subunit plays a crucial role in packaging long strands of DNA into chromosomes via asymmetric loop extrusion [12–14] and ensuring proper individualisation of the chromosomes during cell divisions [15].

**Figure 1.**
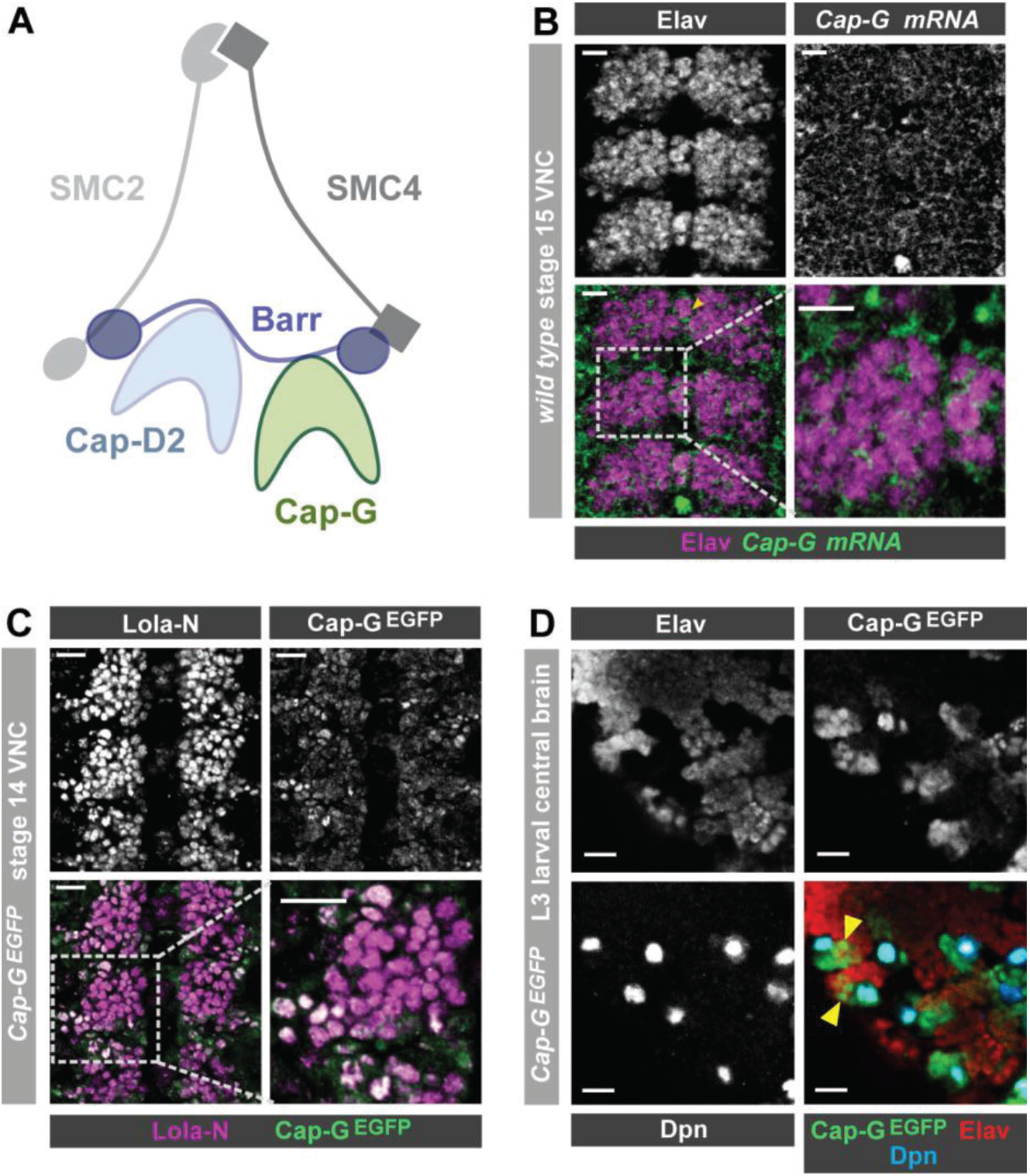
Cap-G expression in *Drosophila* neurons. **A)** Schematic representation of the *Drosophila* condensin I complex. **B)** *w*^*1118*^ embryo (stage 15, anterior top). *Cap-G* mRNA is ubiquitously present in neuronal cytoplasm, neurons marked by Elav. **C)** *Cap-G*^*EGFP*^ embryo (stage 14, anterior top). Cap-G^EGFP^ co-localises with Lola-N in neuronal nuclei of the VNC. **D)** Optic lobe of 3^rd^ instar larvae *Cap-G^EGFP^*. Cap-G is strongly present in NSCs marked by Dpn. Cap-G is present in neurons (Elav positive) in proximity of NSCs (yellow arrowheads). Scale bars 10 μm.

The role of condensin extends beyond its canonical function of physically compacting mitotic chromosomes, with evidence pointing at gene regulation and chromatin organisation [16, 17]. Condensins have been implicated in promoting gene expression by interacting with transcription factors and localising at promoter and enhancer regions. For example, condensin is associated with RNA pol II sites in yeast and mammalian cells [18–20]. Moreover, condensin binding is documented at RNA Pol III (TFIIIC) transcribed regions such as tRNAs in multiple organisms [20–23]. In mouse ESCs, the Cap-H2 subunit of condensin II associates with TFIIIC at highly transcribed, gene-dense regions [23] and when mutated results in reduced gene expression [19]. Condensin I and II are also seen to localise to enhancer regions in human cancer cells, in which they promote transcription through regulation of co-activator recruitment [24]. The role of condensin II in chromatin organisation is evident in *Drosophila*, where Cap-H2 has been associated with regulation of enhancer-promoter interactions [19, 25].

Several gene silencing mechanisms have also been linked to condensin activity. In yeast, chromatin compaction driven by condensin represses transcription in quiescent cells [26]. This is supported by observations in mouse T-cells, where condensin II depletion causes chromatin decompaction and an increase in gene expression, disrupting cellular quiescence [27]. In *Drosophila*, condensin I is implicated in regulation of position effect variegation (PEV) [28–30]. Heterozygous *Cap-G* mutants show wing notches and rough eye phenotypes in flies, which is attributed to a regulatory role of Cap-G in heterochromatin gene expression [28]. Furthermore, the *Drosophila* Cap-H orthologue, Barren, interacts with the chromatin-repressing Polycomb complex to silence homeotic genes [31], whilst condensin II negatively regulates transvection thereby repressing expression of genes at transvection-sensitive loci [32]. In *C. elegans*, the dosage compensation complex (DCC), closely related to condensin I, binds to DNA inducing transcriptional repression [33]. Moreover, depletion of condensin II shows an upregulation in gene expression due to disrupted gene silencing [21]. In murine cells, Cap-G2 has a potential role in erythroid cell differentiation by repressing transcription via chromatin condensation [34]. However, some experiments in yeast argue that condensin does not directly regulate gene expression. For example, a recent study demonstrated that condensin depletion results in genome decompaction but has no effect on overall gene expression [35].

These studies observe the roles of condensin in mitotically proliferating cells, but there is limited knowledge of cell-specific, post-mitotic roles for condensin. Condensin mutations in mitotic and interphase cells ultimately disrupt post-mitotic daughter cells, leading to severe phenotypes. In mouse neuronal stem cells (NSCs), condensin II mutations disrupted nuclear architecture and resulted in apoptosis of NSCs and post-mitotic neurons [36]. Moreover, viable condensin II mutant mice developed T-cell lymphomas and triggered DNA damage in post-mitotic daughter cells [37]. One recent study showed that RNA-levels in *S. cerevisiae* are disrupted upon condensin inactivation due to the well-known phenotype of chromosomal mis-segregation during anaphase [38]. This study points towards condensin having no direct role on transcription and raises the possibility that previous studies implicating condensin in gene expression may be suffering from artefacts resulting from aberrant chromosome segregation. Furthermore, condensin inactivation in differentiated mouse hepatocytes showed no changes in chromatin folding or gene expression [39]. Conversely, a post-mitotic role for condensin II has been demonstrated in *Drosophila*, in which Cap-D3 regulates transcriptional activation of anti-microbial gene clusters in fat body cells [40].

Despite the wealth of knowledge on condensin complexes, their role in post-mitotic cells remains unclear. In this study we reveal a novel role for the condensin I subunit Cap-G in *Drosophila* post-mitotic neurons. We observed Cap-G expression and localisation in the *Drosophila* central nervous system (CNS) *in vivo.* Cell-specific knockdown of Cap-G in neurons resulted in severe developmental arrest, behavioural defects, and an overall misregulation of gene expression in the CNS. Knockdown animals exhibit a downregulation of neuron-specific genes and an ectopic upregulation of non-CNS-specific genes. Finally, Cap-G DNA binding profiles dynamically change between neuronal stem cells (NSCs) and post-mitotic neurons. Cap-G binding is enriched in gene bodies and depleted at accessible chromatin and known enhancer regions. Genes bound by Cap-G significantly overlap with the misregulated genes detected in knockdown animals. The discovery of a neuronal role for Cap-G highlights the importance of studying condensin proteins in a post-mitotic context, to better understand their role in the regulation of gene expression. Furthermore, we show that removal of Cap-G function from immature neurons exhibits a more severe phenotype than when it is removed from more mature neurons, highlighting a key role for condensin in early neuronal development.

## RESULTS

### Cap-G is present in post-mitotic neurons

Upon conducting a yeast-2-hybrid screen to look for proteins interacting with the neuron-specific transcription factor Lola-N, we were surprised to identify the condensin complex component Cap-G as a potential interacting partner. Cap-G is a HEAT-repeat containing subunit exclusive to condensin I [41] (Figure 1A). Whilst condensin activity has been characterised in neural stem cells [36], the role of condensin complexes in post-mitotic neurons has not yet been studied in any species. Published RNA-seq data indicates significant levels of condensin subunit transcripts in neurons, including *Cap-G* [5, 42]. Therefore, we decided to investigate whether Cap-G was indeed present in *Drosophila* post-mitotic neurons. To characterise *Cap-G* expression in the central nervous system (CNS), we carried out fluorescent *in-situ* hybridisation for *Cap-G* mRNA on *w*^*1118*^ embryos (stage 15). We observed ubiquitous expression of *Cap-G* in all cells of the Ventral Nerve Cord (VNC) of *Drosophila* embryos (Figure 1B). *Cap-G* mRNA was clearly detectable in neurons marked by the pan-neuronal marker Elav, indicating post-mitotic expression of *Cap-G* in the CNS. We also observe *Cap-G* expressed broadly in the larval central nervous system in which *Cap-G* mRNA was detectable at similar levels in the neural stem cells (NSCs) and neurons (Figure 1 – figure supplement 1A).

To investigate the distribution of Cap-G protein in the *Drosophila* CNS we utilised a *Cap-G*^*EGFP*^ CRISPR knock-in line. This line expresses C-terminally EGFP-tagged Cap-G from its endogenous locus, is homozygous viable and recapitulates native Cap-G expression and localisation. In stage 14 embryos we observed strong Cap-G^EGFP^ signal in NSCs (Figure 2A) and we were also able to detect nuclear EGFP in post-mitotic neurons, which colocalised with the neuronal transcription factor Lola-N [2] (Figure 1C). As expected, strong GFP signal was observed in mitotic cells in the larval CNS, particularly in NSCs and the optical proliferation centre (Figure 1D, Figure 1 – figure supplement 1B). Cap-G not only co-localised with NSC-marker Deadpan (Dpn), but also with neuronal marker Elav in the larval central brain (Figure 1D). Interestingly, EGFP signal was strongest in Elav-positive neurons in closest proximity to ganglion mother cells (GMCs) and NSCs in both embryos and larvae, indicating that Cap-G may play a more prominent role in newly born neurons. To determine whether Cap-G was the only member of the condensin I complex to be present in post-mitotic neurons, we also examined *barren*^*EGFP*^ animals which express a C-terminally EGFP-tagged Barren variant from their genomic locus. Barren^EGFP^ was clearly detected in all cells of the embryonic VNC as we saw with Cap-G (Figure 1 – figure supplement 1C). Interestingly, we observed that Barren appeared to be predominantly in the cytoplasm of neurons, with much lower signal apparent in the nucleus. This result is in agreement with previous reports which have shown that Barren is primarily localised in the cytoplasm of interphase cells (in which Cap-G appears largely nuclear) [41, 43]. Since Cap-G showed the most distinct nuclear localisation, we decided to focus on the neuronal role of Cap-G for the rest of this study. Overall, these data confirm that condensin expression is prevalent in the post-mitotic cells of the fly CNS and is maintained throughout development, from embryonic to larval stages, indicating that condensin I may have a previously unappreciated role in post-mitotic neurons.

**Figure 2.**
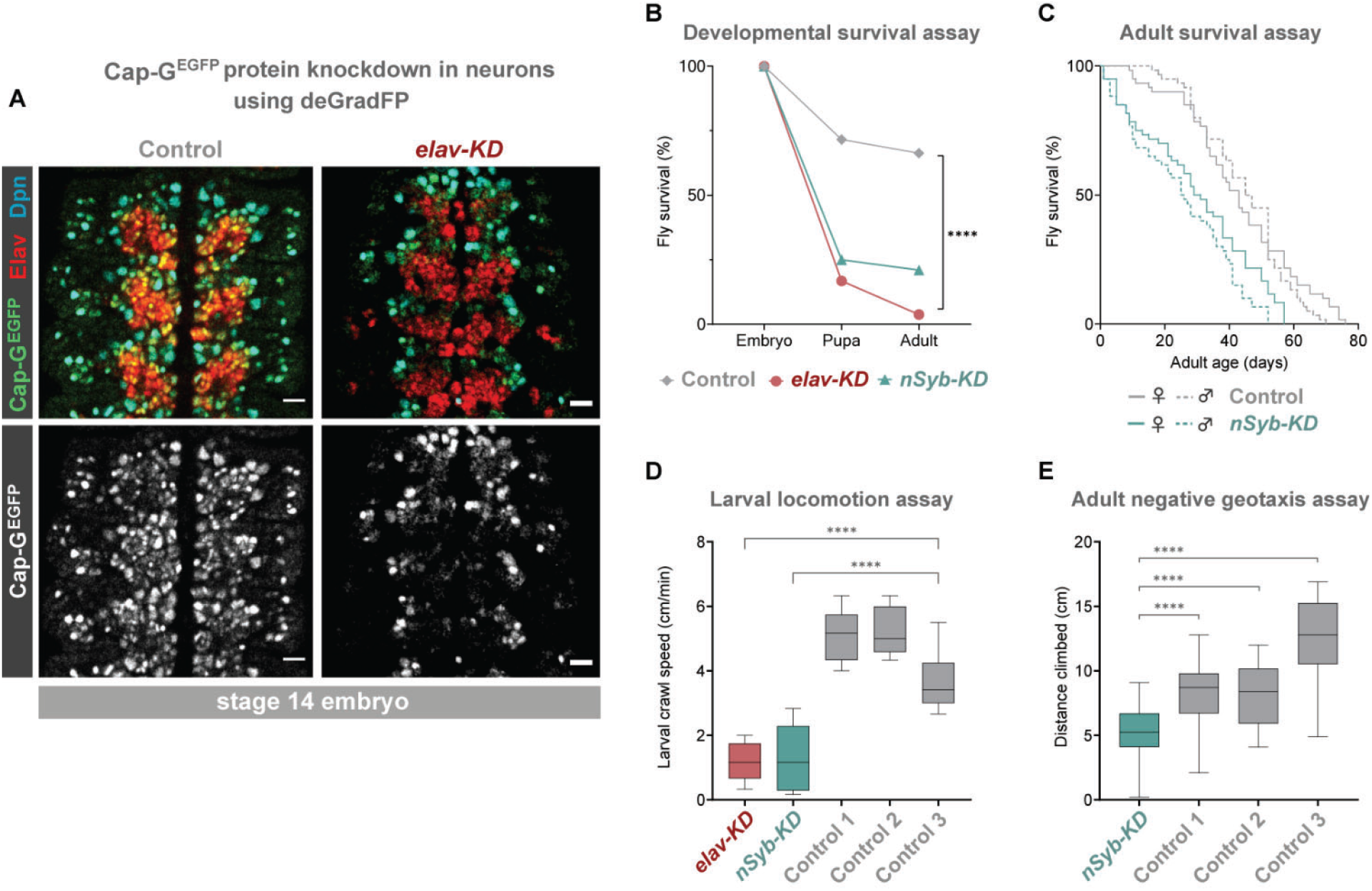
Cap-G knockdown in neurons results in premature developmental arrest and severe mobility defects. **A)** Effective knockdown of Cap-G^EGFP^ in neurons. Embryonic VNC (stage 14, anterior top) showing reduced Cap-G^EGFP^ levels in neurons marked by Elav in *elav-KD* compared to *Cap-G*^*EGFP*^ control. Cap-G^EGFP^ is still detected in NSCs marked by Dpn in both *Cap-G*^*EGFP*^ control and *elav-KD*. **B)** Survival of developing flies recorded at pupal and adult stages. *Cap-G*^*EGFP*^ flies were used as the control genotype. Survival expressed as a percentage of the initial total number of embryos. 300 biological replicates/genotype. Logrank test and weighted Gehan-Breslow-Wilcoxon model (****, p < 0.0001). **C)** Survival of adult *nSyb*-KD and *Cap-G*^*EGFP*^ control recorded daily until all flies deceased. Survival expressed as a percentage of starting flies, 60 biological replicates/genotype. Logrank test and weighted Gehan-Breslow-Wilcoxon model (****, p < 0.0001). **D)** Locomotion assay of L3 larvae from 3 independent experiments. Control 1 = *Cap-G*^*EGFP*^; *UAS-deGradFP*. Control 2 = *elav-Gal4*; *Cap-G*^*EGFP*^. Control 3 = *Cap-G*^*EGFP*^; *nSyb-Gal4*. Mean crawling speed from 10 technical replicates for each of the 30 biological replicates/ genotype. Kruskal-Wallis test one-way ANOVA (****, p < 0.0001). **E)** Negative geotaxis assay from 3 independent experiments. Control 1 = *Cap-G*^*EGFP*^; *nSyb-Gal4*. Control 2 = *Cap-G*^*EGFP*^; *UAS-deGradFP*. Control 3 = *nSyb-Gal4*; *UAS-deGradFP*. Mean distance climbed from 10 technical replicates for each of the 30 biological replicates/genotype. Kruskal-Wallis test one-way ANOVA (****, p < 0.0001).

### The deGradFP system produces robust neuron specific knockdown of Cap-G

The presence of Cap-G in neurons suggests a post-mitotic function for condensin in the CNS. To investigate this putative neuronal role, we used the deGradFP system to create neuron-specific Cap-G knockdown (KD) animals [44]. When combined with *Gal4/UAS*, this system allows for highly specific knockdown of target proteins in a cell-type of interest. By combining deGradFP with *Cap-G*^*EGFP*^ flies, we were able to successfully induce degradation of Cap-G. As expected, ubiquitous degradation of Cap-G-EGFP with *tubulin-Gal4* resulted in 100% embryonic lethality. This is consistent with the essential role of condensin proteins during mitosis as well as previous reports of *Cap-G* mutant embryonic lethality in *Drosophila* [28, 45].

We induced Cap-G knockdown in neurons, using two post-mitotic drivers, *elav-Gal4* and *nSyb-Gal4*. *elav-GAL4* is expressed in all neurons, from newly born to mature, whilst *nSyb-GAL4* drives expression solely in more mature neurons in which synapse formation has begun. We performed knockdown experiments using both *Gal4* lines to better characterise the role of Cap-G during neuronal maturation since we observed higher levels of Cap-G in newly differentiated neurons (Figure 1C, D). Cap-G knockdown experiments with these two drivers are referred to as *elav-KD* and *nSyb-KD* respectively hereafter. Immunostaining of *elav-KD* embryos showed a reduction in GFP signal in neurons but not in NSCs, indicating that a robust neuronal knockdown was achieved using the deGradFP method (Figure 2A). Previous reports have indicated that *elav-GAL4* can drive low level expression in embryonic NSCs (but not post-embryonic NSCs) [46]. Since a recent study raised concerns that condensin knockdown phenotypes seen in interphase may in fact be due to defects in mitotic chromosome segregation [38] we undertook a careful characterisation of the *elav-KD* on NSCs.

To determine whether we could see any reduction in Cap-G^EGFP^ levels in NSCs or GMCs, we quantified fluorescence levels in live mitotic cells of the VNC in *elav-KD* and *Cap-G*^*EGFP*^ control embryos. We observed no difference in Cap-G^EGFP^ fluorescence in mitotic cells between the control and *elav-KD* embryos (Figure 2 – figure supplement 1A) and conclude that any low-level expression of deGradFP in NSCs is not sufficient to significantly reduce Cap-G^EGFP^ levels. Due to the well characterised function of condensin during mitosis, we reasoned that degradation of Cap-G in NSCs would affect their ability to divide. Therefore, we quantified the total number of NSCs (marked by Dpn), the total number of dividing cells (marked by the M-phase marker pH3 [47]) and finally the number of actively dividing NSCs (Dpn and pH3-positive cells). We saw no differences in the overall number of NSCs (Figure 2 – figure supplement 1B), dividing cells (Figure 2 – figure supplement 1C), or number of mitotic NSCs between control and *elav-KD* embryos (Figure 2 – figure supplement 1D).

Depletion of condensin in the mouse cortex resulted in NSC and neuronal apoptosis [36]. Similarly, significant apoptosis was seen in the proliferating cells of zebrafish retinas in Cap-G mutants [48]. Therefore, we sought to determine whether an increased amount of cell death was observable in *elav-KD* NSCs, which could be a result of premature Cap-G depletion. TUNEL staining of embryonic VNCs showed no significant difference in the number of apoptotic cells between *Cap-G*^*EGFP*^ controls and *elav-KD* in either neuronal (marked by Elav) or non-neuronal cells (Figure 2 – figure supplement 2A-C). Together, these data show that *elav-KD* does not significantly lower Cap-G levels in NSCs, it does not affect NSC numbers, nor the number of dividing cells, and does not induce apoptosis. Therefore, we conclude that the *elav-KD* has no effect on NSCs and dividing cells at the embryonic stage, suggesting that with this method we are able to achieve a neuron specific, post-mitotic Cap-G knockdown.

### Neuron-specific knockdown of Cap-G leads to behavioural phenotypes and reduced survival across development

To determine the impact of neuronal depletion of Cap-G on animal survival, we assayed the numbers of *elav-KD* and *nSyb-KD* flies pupating or eclosing as adult flies. We observed a severe developmental lethality phenotype in both neuronal Cap-G KD flies when compared to control *Cap-G*^*EGFP*^ flies. In *elav-KD* animals, only 17% of embryos developed into pupae, and only 4% successfully eclosed to produce adult animals, compared to 71% and 66% reaching pupal or adult stages respectively in controls (Figure 2B). Embryos from *nSyb-KD* flies also displayed a severe survival defect, with only 25% pupating and 21% adults eclosing. Survival analysis using both the Logrank and the Gehan-Breslow-Wilcoxon tests showed a significant difference between the Cap-G^EGFP^ KDs and control flies (*p* < 0.0001, n = 300). Whilst the number of animals surviving to pupation was not significantly different between *elav-KD* and *nSyb-KD,* significantly more *nSyb-*KD flies survived to adult stages. The surviving 4% *elav-KD* adults all died shortly after eclosing.

Despite appearing morphologically normal, we investigated whether surviving *nSyb-KD* flies had any detectable neuronal dysfunction in adulthood. Survival assay for *nSyb-KD* flies revealed a significantly higher mortality rate than controls (Figure 2C). Survival analysis using both the Logrank and the Gehan-Breslow-Wilcoxon tests showed a significant difference between the *nSyb-KD* and control flies (*p* < 0.0001, n = 60). Overall, these results suggest that Cap-G in post-mitotic neurons is necessary for normal development of the CNS and survival.

Further to premature death, we analysed effects of Cap-G KD on larval locomotion ability, commonly used as a proxy to reveal defects in neuronal development [49]. Both *elav-KD* and *nSyb-KD* animals displayed locomotion defects when compared to controls (Figure 2D). On average, the Cap-G knockdown larvae moved 4 cm/min less than the control animals (*p* < 0.0001, n =30). We observed that mobility impairment carried over onto adult flies (Figure 2E). *elav-KD* adults were unable to move after eclosion and died shortly after. A negative geotaxis (climbing) assay [50] on *nSyb-KD* adults showed a compromised ability to climb, with the mean climbing distance of ~5 cm, as compared to ~9 cm or ~13 cm for control genotypes (Kruskal-Wallis test one-way ANOVA, p < 0.0001, n = 30) (Figure 2E). Overall, these data show that knockdown of Cap-G in neurons results in motor defects in both larvae and adult flies.

### Neurons have increased levels of DNA damage due to post-mitotic Cap-G KD

We reasoned that due to the role of condensin complexes in organising chromosomes, depletion of Cap-G may have resulted in genomic instability, therefore we decided to assay DNA damage in Cap-G knockdown flies. Using a ɣ-H2Av antibody as a marker for DNA damage [51] and Elav as a neuronal marker, we quantified the percentage of ɣ-H2Av positive neurons in larval brains. In the larval CNS we observed an increase in neuronal DNA damage in both *elav-KD* and *nSyb-KD*, when compared to *Cap-G*^*EGFP*^ controls (Figure 3A). In *elav-KD* and *nSyb-KD* we observed a two/threefold increase of ɣ-H2Av+ neurons compared to controls, (Kruskal-Wallis test one-way ANOVA, p < 0.001) (Figure 3B).

**Figure 3.**
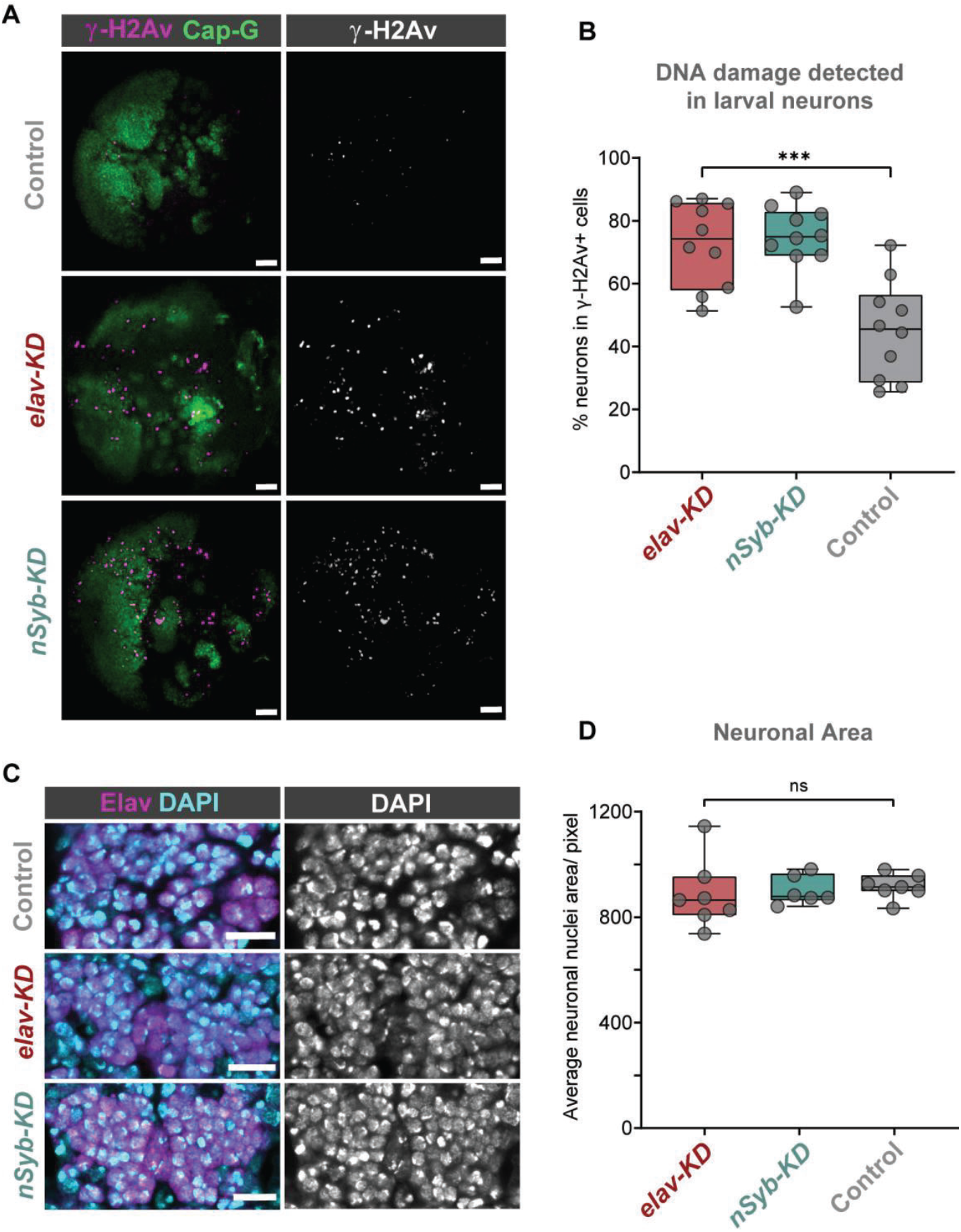
Cap-G knockdown leads to increased DNA damage in larval neurons. **A)** Larval optic lobes for *Cap-G*^*EGFP*^ controls and *Cap-G KD* genotypes. ɣ-H2Av indicates DNA damage, Cap-G signal is strongest in dividing cells. Scale bars 10 μm. **B)** Quantification of ɣ-H2Av+ neurons as percentage of total ɣ-H2Av+ cells. 10 biological replicates per genotype. Kruskal-Wallis test one-way ANOVA (***, p < 0.001). **C)** Section of embryonic VNC (ventral view) stage 15 for all genotypes. Elav to mark neuronal nuclei and DAPI as nuclei stain. Control (*Cap-G^EGFP^*), *elav-KD* and *nSyb-KD* genotypes. **D)** Quantification of neuronal nuclei area. Pixel area average of at least 1000 cells per replicate was analysed. 7 biological replicates per genotype. No statistical difference determined by one-way ANOVA (p> 0.5).

Given that condensin complexes are implicated in chromatin condensation, it seemed reasonable to question whether Cap-G has any role in regulating chromatin structure or distribution in neurons. To assess whether Cap-G KD influenced overall chromatin condensation we quantified nuclear area in embryonic neurons. Control and KD embryos were stained for the neuronal marker Elav as well as DAPI to highlight overall DNA content (Figure 3C). We observed no significant difference in the size of nuclear area between Cap-G KD and controls when Cap-G was depleted with either *elav* or *nSyb-Gal4* (Figure 3D).

### Dynamic Cap-G association with chromatin in mitotic NSCs and post-mitotic neurons

Since we have observed Cap-G in the nuclei of post-mitotic neurons, we sought to characterise its association with chromatin to further illuminate its role in gene regulation in the CNS. We used Targeted DamID (TaDa) [52, 53] to profile cell-type specific Cap-G binding in NSCs and post-mitotic, differentiated neurons. For differentiated neurons, we continued to use both *elav* and *nSyb-Gal4* drivers to further characterise differences in Cap-G binding between newly born and fully differentiated neurons (Figure 4A). Cap-G binding in NSCs was determined using *wor-Gal4* so that we could compare the genomic localisation of Cap-G between actively dividing and post-mitotic cells. TaDa requires the expression of a fusion protein with the *E. coli* Dam methylase. Since we have seen that Cap-G remains functional when tagged with EGFP at the C-terminus, the fusion of Dam at the same site should not disrupt Cap-G localisation or function.

**Figure 4.**
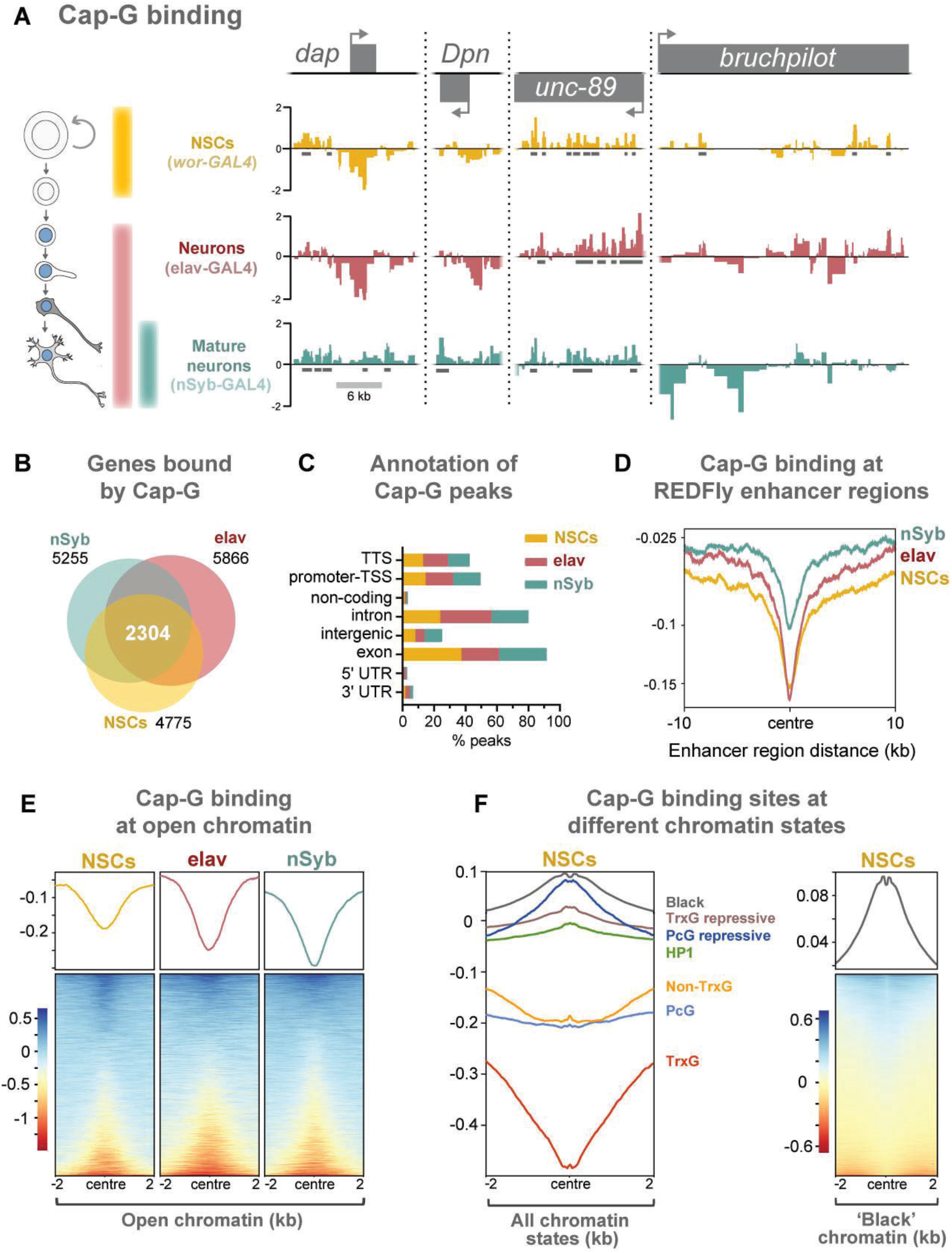
Cap-G binds to DNA in NSCs and neurons. **A)** Cap-G binding at example genes in NSCs (*wor-Gal4*), all neurons including immature neurons (*elav-Gal4*) and mature (*nSyb-Gal4*) neurons. Horizontal grey bars indicate statistically significant peaks. **B)** Venn diagram showing unique genes bound by Cap-G and the total overlap between all cell types. **C)** Genomic annotation of Cap-G peaks shows enrichment in gene bodies (introns and exons) whilst binding at non-coding regions are depleted. **D)** Cap-G binding is depleted at known Drosophila Regulatory Elements (REDfly). Profiles plotted against centre of enhancer region and 10 Kb up/downstream. **E)** Average Cap-G binding is depleted at accessible chromatin for all cell types. Plot shows 2 Kb up/downstream centre of an open chromatin region. **F)** Average Cap-G binding in different chromatin states in NSCs. Cap-G binding is enriched in repressive states (Black, HP1, TrxG-repressive, and PcG repressive), and depleted in permissive chromatin states (non-TrxG, TrxG, and PcG). Heatmap is shown for most highly enriched (black) state.

The DamID methyl-PCR required for the TaDa experiments yielded strong amplification for all cell types, indicating that Cap-G-Dam associates with chromatin in post-mitotic cells. Sequencing revealed that Cap-G peaks are prevalent across the genome of all cell types. Furthermore, Cap-G binding is highly variable between NSCs and differentiated neurons, suggesting specific roles in gene regulation in these cell types (Figure 4A, Figure 4 – figure supplement 1). Cap-G binds to known NSC-specific genes such as *deadpan* and *dacapo* in mature neurons, but not in NSCs, where these genes are actively expressed [54, 55] (Figure 4A). Conversely, Cap-G peaks are detected at the neuronal gene *bruchpilot* in NSCs but are absent in nSyb neurons. Moreover, Cap-G also binds to non-neuronal genes in all cell-types, such as *unc-89*, which is expressed in muscle tissues, but not the CNS [42]. Overall, the number of genes bound by Cap-G is similar across cell types, but significant differences are apparent with uniquely bound genes detected in each cell-type (Figure 4B). Interestingly, marked differences are apparent in Cap-G binding between the two post-mitotic cell types. Together these data suggest that Cap-G binding is cell-specific and dynamic across cell types, changing as the NSC differentiates and the neuron matures.

Cap-G peaks are strongly enriched in gene bodies and depleted in intergenic and non-coding regions (Figure 4C). We did not observe significant binding of Cap-G at tRNA genes or rRNA genes in our data, as previously reported for condensin binding in *S. cerevisiae* (in which there is only one condensin complex) or condensin II binding in mouse ESCs [20, 23].

### Chromatin accessibility is reduced at Cap-G binding sites across cell types

Previous evidence suggests that condensins associate with open chromatin and enhancer regions in *C. elegans* [21]. Similarly, the condensin II subunit, Cap-H2 localises at enhancer regions in *Drosophila* Kc167 cells [25]. We analysed the Cap-G binding at known Regulatory Elements in *Drosophila* [56] and observed a strong depletion of Cap-G binding at those sites, suggesting that Cap-G is not strongly associated with enhancers in the *Drosophila* CNS (Figure 4D). The recently described CATaDa method [9] enabled us to use the control Dam-only binding from all three cell types as a reference for accessible chromatin and compare it to Cap-G binding. Overall open chromatin regions display a depletion of Cap-G binding (Figure 4E). We also observed a negative correlation between chromatin accessibility and Cap-G binding across all cell types (Figure 4 – figure supplement 1A, B).

It has been demonstrated that chromatin can be divided into several discrete states depending on the occupancy of various key proteins [57]. These states include repressive and permissive chromatin environments. Recently published data allowed us to compare Cap-G binding to chromatin states in NSCs [58]. We find that Cap-G binding is most strongly enriched in repressive chromatin states (Figure 4F). Cap-G was particularly strongly associated with ‘black’ chromatin – a prevalent repressive state that does not incorporate traditional heterochromatin markers. In contrast, Cap-G binding was relatively depleted at permissive chromatin states, particularly the ‘red’ TrxG state (Figure 4F).

Overall, our results suggest that accessible chromatin is depleted at Cap-G binding sites indicating that condensin I does not bind to known enhancer and regulatory regions in *Drosophila* NSCs and neurons. Together these data suggest that Cap-G binding of chromatin in neurons may have a regulatory role independent of binding to accessible chromatin and may promote, or be recruited to, repressive chromatin environments.

### Knockdown of Cap-G in neurons leads to misregulation of gene expression in the CNS

Having observed Cap-G in neurons and with specific DNA binding properties in differentiated cells led us to further investigate the role of Cap-G in neuronal gene regulation. We performed RNA-seq experiments on the CNS of neuron-specific Cap-G knockdown flies. Due to the differences we had previously observed between *elav-KD* and *nSyb-KD* phenotypes, we continued to use both drivers to investigate differences in gene expression when Cap-G is depleted either in newly born and mature neurons (*elav-KD*), or more mature neurons (*nSyb-KD*). Cap-G knockdown resulted in disruption of the neuronal transcriptome, with a significant number of genes differentially expressed using both *elav* and *nSyb* drivers (Figure 5A, Figure 5 – figure supplement 1A, B). *elav-KD* resulted in 1360 upregulated and 1308 downregulated genes, whilst in the *nSyb-KD* we observed a more modest effect on gene expression, with 152 upregulated and 126 downregulated genes. Of genes that are differentially expressed in both experiments (83 genes), a similar pattern of up/downregulation was observed (Figure 5D).

**Figure 5.**
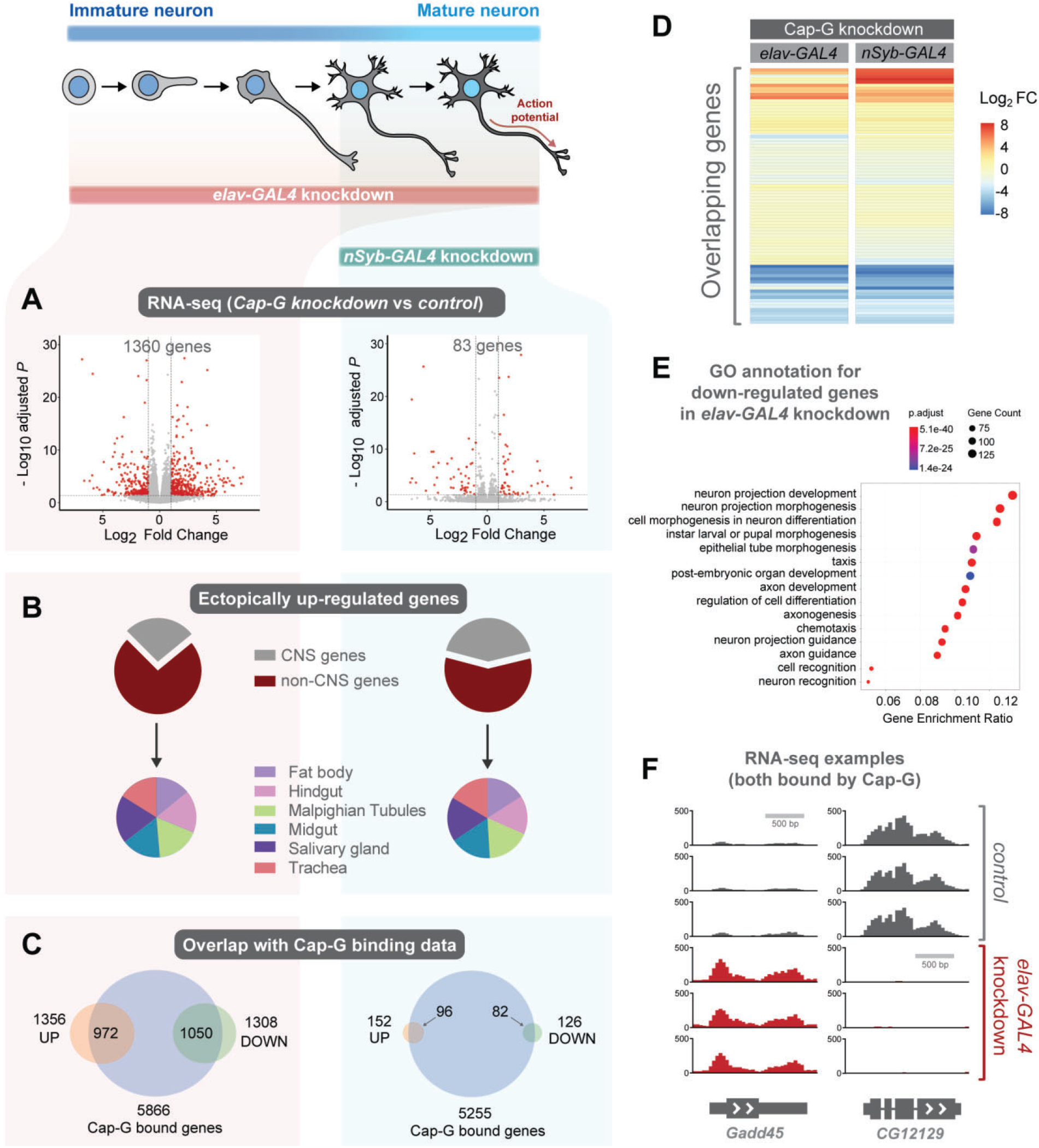
Cap-G knockdown in neurons display severe misregulation of gene expression. **A)** Volcano plots showing differentially expressed genes in *elav-KD* and *nSyb-KD* respectively. Significant differential expression marked in red by a Log2 fold change >1 or <-1 and FDR <0.05. Control genotypes are *elav-Gal4; UAS-deGradFP* and *nSyb-Gal4; UAS-deGradFP* for *elav-KD* and *nSyb-KD* respectively. **B)** Pie charts showing tissue of origin expression data for upregulated genes in Cap-G KD datasets taken from FlyAtlas. The majority of genes are not normally expressed in the CNS but in alternative tissues. **C)** Venn diagrams showing significant overlap between Up / Down regulated genes and Cap-G DamID peaks for elav and nSyb neurons respectively, Fisher’s exact test (p < 1× 10^−20^). **D)** Heatmap of overlapping differentially expressed genes between *elav-KD* and *nSyb-KD*. **E)** Enriched Gene Ontology analysis of downregulated genes from *elav-KD* shows mostly neuron-specific terms. **F)** Example of differentially expressed genes directly bound by Cap-G in *elav-KD* (all three replicates displayed).

Analysis of Enriched Gene Ontology (GO) terms shows that downregulated genes in *elav-KD* were tissue-specific, with neuron-specific terms such as “neuron projection development” and “cell morphogenesis in neuron differentiation” being highly enriched (Figure 5E). Gene expression data extracted from FlyAtlas 2 [42] revealed that the majority of upregulated genes are not normally expressed in the larval CNS (Figure 5B). 73% of upregulated genes in *elav-KD*, and 58% in *nSyb-KD* are non-CNS specific, belonging to tissues such as the Midgut, Hindgut and Fat body. Interestingly, the tissue distribution of these ectopically expressed genes is very similar in both knockdown scenarios. This is further confirmed by GO term analysis of upregulated genes that yielded a variety of non-CNS specific GO terms for both knockdowns, such as “nucleotide metabolic process” and “midgut development” (Figure 5 – figure supplement 1C, D). These data suggest that Cap-G contributes to the repression of non-neuronal gene expression following terminal differentiation.

To determine whether misregulation of gene expression was a direct consequence of loss of Cap-G binding in the genome we compared our RNA-seq data to the previously described Cap-G binding profiles (Figure 4). We observed a significant overlap between Cap-G bound genes in our TaDa data and the differentially expressed genes for both *elav* and *nSyb-KD* (Figure 5C, F). In *elav* expressing neurons, 80% of downregulated genes and 72% of upregulated genes were directly bound by Cap-G. These overlaps are highly significant (Fisher’s exact test, p < 10^−20^). In mature neurons, 63% of upregulated genes and 65% of downregulated genes were bound by Cap-G (Fisher’s exact test, p < 10^−20^). These data suggest that the majority of misregulated gene expression observed may be a result of direct binding by Cap-G.

## Discussion

Cell differentiation during development and tissue regeneration requires extensive transcriptional changes with concomitant chromatin remodelling. This has been extensively studied with respect to the differentiation of stem cells and progenitor cells towards a post-mitotic state [59]. However, the alterations in transcription and chromatin during maturation of post-mitotic cells have been relatively unstudied. We have previously shown gradual changes in chromatin accessibility during neuronal maturation [9], including a loss of accessibility at neural stem cell genes. The genomic binding of key chromatin modifiers is also dynamic during this process [58]. Furthermore, promoter accessibility and the expression of axonal regeneration genes decrease in maturing retinal ganglion cells [60, 61]. These gradual changes of the chromatin state could reflect a steady change in cell fate plasticity. This is supported by the discovery that neurons lacking specific transcription factors can dedifferentiate [2, 62] and that the ability of neurons to regenerate axons is dependent on their epigenetic state [63, 64]. Although this field is in its infancy, emerging methods are poised to further advance our understanding [59]. Here, we have discovered a role for the condensin I subunit Cap-G in neuronal development, where its function is particularly important during the early stages of differentiation.

Whilst condensin function in chromosome segregation is well defined, non-canonical roles for condensin complexes in the regulation of gene expression are less well understood. Despite this, recent studies have provided evidence of condensin proteins contributing to gene regulation and chromatin organisation [65]. Whilst it seems certain that condensin complexes play a role in gene regulation, some of these studies may suffer from technical artefacts relating to premature condensin knockdown in dividing cells [38]. Furthermore, studies examining condensin function outside of mitosis have drawn conflicting conclusions [39, 40]. In this manuscript we aimed to characterise the role of a condensin I protein (Cap-G) in a previously unstudied post-mitotic cell type. We found that Cap-G plays a significant part in regulating gene expression in neurons which is necessary for normal nervous system function.

We observed nuclear localisation of Cap-G in *Drosophila* NSCs and neurons during development. The presence of *Drosophila* Cap-G in the nuclei of post-mitotic neurons mirrors its reported nuclear localisation of Cap-G in *Drosophila* interphase of mitotically proliferating cells [41]. Previous analyses of condensins in the central nervous system showed that condensin I and II are both involved in murine NSC division, but “largely absent” from neurons [36]. It is possible that the relative intensity of signal observed from dividing and post-mitotic cells masked the presence of condensin subunits in this instance. However, both our *in situ* hybridisation and fluorescently tagged protein imaging, as well as previously published RNA-seq data, strongly suggest that Cap-G is prevalent in *Drosophila* post-mitotic neurons. Mammalian neurons may not utilise Cap-G in post-mitotic neurons to regulate gene expression. Further studies will be required to determine whether this property is conserved.

To understand the role of Cap-G in the CNS we induced neuron-specific KD which resulted in premature developmental arrest and severe mobility impairment of flies (Figure 2). *Cap-G* mutants in *Drosophila* are homozygous lethal, however heterozygous animals for some alleles showed rough eyes phenotypes [28]. Since we did not observe any eye phenotype in our neuron specific knockdowns (which include eye tissue), we surmise that the rough eye previously reported in Cap-G heterozygotes is due to misegregation or chromatin organisation defects arising from knockdown in progenitor cell-types. This suggests that the phenotypes we observe in neuronal knockdowns may be a consequence of a separate post-mitotic role for Cap-G.

We depleted Cap-G levels in two overlapping populations of cells, fully differentiated neurons (expressing *nSyb*), as well as immature to fully differentiated neurons (expressing *elav*). *elav-Gal4*-driven knockdowns consistently displayed more severe phenotypes than with *nSyb-Gal4* and had correspondingly more drastic changes in gene expression. This difference in phenotype can be attributed to the cells targeted by the different drivers. e*lav-Gal4* encompasses all neuronal cells, including newly born neurons that are in a more transient and plastic state, when compared to more mature, synapse forming-neurons targeted with *nSyb-Gal4*. This hints that Cap-G may serve a role in terminal differentiation of newly born neurons, but also maintenance of neuronal cell-state once matured. Consistent with this we saw much higher levels of Cap-G in newly born neurons than more differentiated neurons, suggesting that Cap-G may play a more prominent role in the early life of neurons. Previous reports have stated that *elav-Gal4* may drive expression prematurely in embryonic NSCs [46]. We made extensive efforts to determine whether *elav-Gal4* caused a reduction in Cap-G levels in NSCs or any defects in NSC division. Since we could not see any changes in Cap-G levels in NSCs and did not observe any phenotypes that we would expect to see from mitotic Cap-G knockdown, we concluded that the phenotypes we observed were a result of post-mitotic Cap-G depletion in neurons. That phenotypes were observed with two independent Gal4 lines provides further confidence in a *bona fide* neuronal role for Cap-G. In future, the development of Gal4 lines with even more precise spatial and temporal control will allow us to carefully interrogate the differences between condensin function in the different stages of neuronal maturation.

We initially decided to investigate Cap-G function in neurons based on a potential interaction with the neuronal transcription factor Lola-N. Lola_N is required in the early stages of neuronal differentiation to maintain the differentiated cell state [2]. Similarly, we observed increased levels of Cap-G and more severe consequences from knockdown of Cap-G in the early stages of neuronal maturation. Therefore, it is intriguing to speculate that Cap-G/condensin I act together with Lola-N to maintain the neuronal transcriptome in differentiating neurons. Further studies will be necessary to verify whether there is in fact a functional relationship between these two proteins.

Our data suggest that DNA damage is significantly increased in Cap-G depleted neurons. Previous studies in yeast have implicated the condensin complex in the regulation of the DNA damage response during interphase [66, 67]. In human cells condensin I was shown to have a function in single stranded break (SSB) specific DNA damage repair but was not involved in double stranded break (DSB) repair [68]. In contrast, the antibody used in our DNA damage experiments specifically recognises the response to DSBs. Condensin proteins have not yet been shown to function in DNA damage pathways in *Drosophila*, therefore it is possible that condensin I is involved in DSB repair in flies, in contrast to mammalian cells. However, Cap-G/condensin I may be required to maintain genomic stability in neurons independently of the DNA damage response. It is unclear whether the DNA damage we observe is responsible for the changes in gene expression. Given that we see similar levels of DNA damage in neurons of both *elav* and *nSyb* mediated knockdown, which have significantly different levels of aberrant gene expression, it is likely that increased DNA damage is not directly responsible for the changes we see in gene expression.

Previous studies have demonstrated condensin binding at genes the expression of which are thought to be directly regulated by association with condensin. We demonstrated that Cap-G binds dynamically to DNA in NSCs and post-mitotic neurons. Each cell type displayed unique genes bound by Cap-G. Interestingly, Cap-G peaks are enriched mid-gene and not at promoter regions as reported in *C. elegans* and chicken DT40 cells [21, 22]. Moreover, we did not observe any overlap between Cap-G binding and TFIIIC targets, as previously described in multiple species [20–23]. Condensin binding data reported to date has been collected from mitotic or interphase cells. Therefore, condensin binding patterns in terminally differentiated cells are unknown, and the binding patterns we observe may be unique to this cell type. Similarly, we saw that Cap-G binding was depleted at accessible chromatin and known enhancer and regulatory element regions [42]. This is contrary to a study in which Cap-G was shown to localise to active enhancers in human cancerous cells, promoting estrogen-dependent gene expression [24]. However, conflicting evidence for condensin I binding at enhancer regions has been presented. For example, Barren has been shown to overlap with few known enhancers in *Drosophila* Kc167 cells [69].

Our results show that Cap-G depletion leads to alterations in gene expression in both early and mature neurons in *Drosophila* larvae. We observed that neuron-specific genes are downregulated, whilst non-CNS genes are ectopically upregulated in neurons depleted of Cap-G, suggesting that Cap-G activity contributes to maintenance of the neuronal transcriptome. These observations are supported by previous studies in interphase and mitotic cells which show that Condensin I and II regulate cell-specific gene expression in multiple species [23, 24, 27]. However, only a single study reports a post-mitotic role for condensin, specifically condensin II subunit Cap-D3, in *Drosophila* [40]. The authors describe Cap-D3 regulating cell-specific gene expression in fat body cells, where it binds to and transcriptionally activates innate-immunity genes, including antimicrobial peptides. Interestingly, adult flies depleted of Cap-D3 have impaired immune response and fail to effectively clear bacteria [40]. Therefore, condensin I and II subunits may have different roles according to developmental stages, cell types, and complex incorporation.

To date, the role of condensin in transcription regulation has only been speculated for interphase and dividing cells. For example, condensin involvement in transcriptional repression has been suggested in *C. elegans* [21], mouse T-cells [27] and yeast [26]. However, there is conflicting evidence in the literature on the role of condensin in gene expression. A recent study in fission and budding yeast suggests that condensin has no direct effect on gene expression and that changes to RNA levels occur as an indirect effect of chromosome mis-segregation in condensin-depleted cells [38]. Moreover, recent studies in yeast and mouse hepatocytes showed no significant changes in gene expression upon condensin depletion [35, 39]. Our data is exclusively post-mitotic therefore there is no effect on chromosome stability, emphasising that the effects we see on neuronal gene expression are a direct result of Cap-G depletion. Furthermore, a large proportion of the affected loci appear to be associated with Cap-G, suggesting that Cap-G could have a direct transcriptional regulatory effect on its binding regions. This is also observed in *C. elegans*, where the dosage compensation complex binds to specific DNA regions and promotes transcriptional repression [33, 70].

The canonical role of condensin is as a molecular motor that extrudes DNA loops thereby organising chromosome architecture during mitotic chromosome compaction [71]. Given that condensin has this ability to rearrange chromosome topologies, it is feasible that the action of Cap-G in neurons could be to remodel 3D chromatin structure for optimal gene expression. Since we see more severe consequences of Cap-G depletion in younger neurons, this may suggest that condensin is required to establish a mature post-mitotic chromosome conformation in fully differentiated neurons before synaptogenesis. Another SMC complex, cohesin, is well known to be required for the formation of topologically associated domains [10]. Perturbation of cohesin in *Drosophila* neurons resulted in behavioural phenotypes and defects in axon pruning, also indicating that remodelling of chromatin architecture is likely to be important in neuronal maturation [72].

In conclusion, we have shown that neuronal Cap-G is required for normal development and survival in *Drosophila.* This implicates the condensin complex in a previously uncharacterised role in gene expression in post-mitotic cells. Further studies will be necessary to fully define the mechanism by which this regulation is mediated, whether that is by direct regulation of gene expression, or indirectly, through remodelling the topology of the 3D genome.

## Materials and Methods

### Fly stocks

*Cap-G*^*EGFP*^ and *barren*^*EGFP*^ CRISPR knock-in lines were generated by first inserting cassettes directing expression of EGFP in the eye, immediately downstream of the *Cap-G* and *barren* reading frames within the context of their genomic loci. This facilitated easy screening of knock-in individuals due to their green eye fluorescence. Upon FLP-recombinase mediated excision of the eye-specific promotor, eye fluorescence was lost and at the same time a continuous reading frame was generated between *Cap-G* or *barren* and *EGFP*. Thus, C-terminally EGFP-fused Cap-G and Barren variants are expressed under control of their genomic regulatory elements. Details of the strain construction will be described elsewhere. For degradFP experiments we used the line *w*;; P[w^+mC^=UAS-Nslmb-vhhGFP4]* [44]. *tubulin-Gal4/TM6B* was used for ubiquitous degradation of Cap-G. *UAS-mcd8-GFP* was used for fluorescent reporter experiments. Neuron-specific Cap-G knockdown was driven by the following GAL4 lines *P(GawB)elav^C155^-GAL4* for newly born neurons and *w^1118^; P{y^+t7.7^ w^+mC^=GMR57C10-GAL4}attP2* (*nSyb-GAL4* - Bloomington #39171) for mature neurons. *wor-Gal4* was used to drive expression in NSCs [73].

### Immunohistochemistry and in-situ hybridisation

Third instar larvae and adult CNS were dissected in 1x PBS and tissue was fixed for 20 minutes in 4% formaldehyde (Polysciences, Inc., 10% methanol-free) diluted in PBST (0.3% Triton X-100 in PBS). Tissue washes were done in PBST every 5-15 minutes. Normal Goat Serum (2 % in PBST) was used for tissue blocking (RT, 15 min-1 hour) and subsequent overnight primary antibody incubation. Tissue was mounted on standard glass slides in Vectashield Mounting medium (Vector laboratory).

Embryos were kept at 25 °C and collected every 12-15 hours. Embryos were prepared for confocal imaging using a standard protocol as previously described [2]. Embryos were dechorionated using 50% bleach solution and subsequently fixed in a 1:1 solution of 4% formaldehyde to Heptane. After fixation embryos were washed in 100% methanol to ensure removal of the vitelline membrane and subsequently washed with PBST. Tissue was blocked in 2% NGS (RT, 15 min-1 hour) and then incubated with appropriate primary antibodies overnight.

Primary antibodies used include: rat anti-elav 1:500 (Developmental Studies Hybridoma Bank, DSHB), chicken anti-GFP 1:2000 (Abcam), rabbit anti-phospho-Histone H3 Ser10 (pH3) 1:500 (Merck Millipore, 06-570), guinea pig anti-deadpan 1: 1000 (provided by A. Brand), rabbit anti-Lola-N 1:10 [2] and mouse anti-ɣ-H2Av 1:500 (DSHB - [51]). Secondary antibodies used include Alexa Fluor® 488, 545 and 633 1:200 (Life technologies) and tissue was incubated for 1.5 hours at room temperature. The DNA stain DAPI (1: 20,000) was used as nuclear counterstain.

TUNEL staining on embryos was performed using the Click-iT® TUNEL Alexa Fluor® 549 (ThermoFisher). The provided protocol was followed. The only optimisation included incubation of fixed embryos with Proteinase K for 30 minutes at room temperature and subsequent post-fixation in 4% formaldehyde for 15 minutes at room temperature.

For *in-situ* hybridisation experiments, RNA probes were designed using LGC Biosearch Technologies’ Stellaris® RNA FISH Probe Designer against *Cap-G* exon 4 and with Quasar® 570 dye. Tissue was treated following the published protocol [74]. Fixed and blocked tissue was incubated in hybridisation buffer with *Cap-G* probe (3μM) and primary antibody of interest for 8-15 hours.

### Imaging and Image Analysis

Samples were imaged using a confocal microscope Zeiss LSM 510 and Leica CF8. Analysis of acquired images was done using the Fiji [75] and Icy-Bioimage Analysis software [76]. Icy plugin Spot Detector [77] was used to analyse the total number of cells when quantifying dividing NSCs, pH3 staining, DNA damage (ɣ-H2Av) as well as TUNEL staining, filters were adjusted per experiment and kept constant across control and experimental images. For nuclear area analysis the Ilastik software was used for bulk image segmentation, [78] to recognise each nucleus as an individual ROI. Subsequently, the segmented image was analysed in Fiji using Analyze Particles and extracting the mean nuclei area (>1000 cells / replicate) in a total of 7 embryonic replicates. To quantify live GFP levels we took a z-stack (4 slices minimum of the VNC) per embryo. Actively dividing cells displayed highest GFP levels, therefore binary images were created by using a threshold (Outsu’s thresholding) to select dividing cells as regions of interest (ROI). Mean pixel intensity per ROI (dividing cell) was calculated using Analyze Particles plugin in Fiji (> 30 cells/ embryo). Average pixel intensity of ROI was calculated across z-stack slices, for a total of 10 biological replicates.

### Behavioural and phenotypic assays

All animals were kept at 25 °C unless otherwise stated. For the developmental survival assay three biological replicates per genotype were used. Flies were allowed to lay for 5 hours and then 100 embryos per replicate were collected. After 24 hours, L1 larvae were transferred to a food vial and allowed to develop. The number of pupae (5 days post- lay) and eclosed adults (10 days post-laying) were recorded. Adult survival assays were performed on three replicates, with 20 flies per replicate. Male and female animals were separated upon eclosion. Death was scored daily until total number of flies deceased.

Behavioural assays were performed as previously described [79]. Locomotion assay was performed on 3^rd^ instar larvae using 10 biological replicates and 3 technical replicates. Individual larvae were setup in clear agar plates against a 0.5 cm^2^ grid. Larvae were left to acclimatise for 1 minute and locomotion was recorded as distance covered per minute. Negative geotaxis assays were performed on 10 male flies and 10 technical replicates per genotype. Flies were set up in climbing vials against a grid and a camera. Video recording was started before sharply tapping flies to the bottom of vials. Videos were analysed using Icy, the frame in which the first fly reached the top of the vial was extracted. The coordinates of each fly were used to calculate the average distance climbed per replicate. Statistical tests and plots were performed using the software GraphPad Prism version 8 for Windows.

### Targeted DamID

We generated a *UAS-Cap-G-Dam* line to use in Targeted DamID experiments. The isoform Cap-G-PF was amplified by PCR from cDNA library. The *Dam* construct was fused on the C-terminus of Cap-G using fusion PCR. The *Dam* sequence is frequently fused to the N-terminus of the protein of interest, however, the fusion of a protein to the Cap-G C-terminus has been previously shown to have no effect on protein integrity or function [41]. The *mCherry* sequence used for the primary ORF in the TaDa cassette was amplified from pUAST-LT3-Dam [52] template and fused by PCR to the N-terminal of *Cap-G-Dam* construct. Finally the *mCherry-Cap-G-Dam* construct is cloned into the pUAST-attB [80] plasmid using Gibson Assembly [81].

To infer cell-specificity in our Targeted DamID [52] experiments we used the following driver lines: *wor-Gal4; tub-Gal80ts* for neuronal stem cells, *elav-Gal4; tub-GAL80ts* for immature and mature neurons and *tub-GAL80ts; nSyb-Gal4* for mature neurons. The *UAS-Cap-G-Dam* line was used to profile Cap-G binding and the Dam-only line *tub-GAL80ts; UAS-LT3-NDam* [52] was used as a control for Dam expression and Chromatin Accessibility TaDa experiments [9].

Animals were crossed to the desired driver line and embryos collected at 25°C for 4 hours. To obtain third instar larval brains, embryos were raised at 18 °C for seven days post collection and subsequently placed at 29 °C for 24 hrs to induce Dam- expression. 30 larval brain per replicate were dissected in PBS with 100mM EDTA. Extraction of Dam-methylated DNA and genomic libraries were performed as previously described [82]. Illumina HiSeq single-end 50 bp sequencing was performed on two replicates. Sequencing obtained was normalised against *Dam-only* control binding and mapped against the *Drosophila* genome (r6.22) as described in the published protocol [82].

### Peak calling and annotation

Significant peaks in the Cap-G binding profile were analysed using a previously described DamID peak-calling pipeline [82]. A false discovery rate (FDR) is calculated and assigned to all potential peaks. Those with an FDR less than 1% were classified as significant. Any gene identified within 5 kb of a peak (with no interfering genes) was classified as a potentially regulated gene. Cap-G peaks were annotated using HOMER software annotatePeaks pipeline [83].

### RNA-seq and data analysis

RNA-seq was performed on three biological replicates, for Cap-G knockdown and controls respectively. Control genotypes were *elav-Gal4; UAS-deGradFP* and *nSyb-Gal4; UAS-deGradFP* for *elav-KD* and *nSyb-KD* respectively. Total RNA was extracted from dissected 3^rd^ instar larvae CNS, 35 per replicate, using a standard TRIzol ® extraction protocol [84]. RNA library preparation and sequencing was performed by Beijing Genomics Institute (BGI).

Sequencing data was mapped to the *Drosophila* genome (release 6.22) using STAR [85]. Mapped files were collected in a matrix using *featureCounts* from the Rsubread package [86]. Differential expression analysis and MA plots were carried out using the Deseq2 R package [87]. Volcano plots were generated using EnhancedVolcano R package [88]. Genes that had an adjusted p-value <0.05 and a log_2_ fold change greater than 1 (for upregulated) or less than −1 (for downregulated) were classified as significant. Heatmap generated using pheatmap package in R.

Tissue expression data from FlyAtlas 2 was used to determine tissue of origin of upregulated genes [42]. Larval FPKM values above 2 were considered non-background and were used as a threshold for tissue specificity. Expression data for a list of genes of interest was extracted from the FlyAtlas 2 database and data analysis carried out using Pandas in Python [89].

### Statistical analysis and Gene Ontology

Overlap analysis of peak-files from different datasets was done using Bedtools Intersect and statistical significance determined by Fisher’s exact test using Bedtools fisher [90]. Deeptools was used to generate average signal profiles, heatmaps, principal component analysis and correlation matrices [91]. Enrichment GO analysis was performed on gene lists of interest using the R package clusterProfiler [92]. All other figures were produced using the ggplot2 package in R.

### Data availability

All raw sequence files and processed files have been deposited in the National Center for Biotechnology Information Gene Expression Omnibus (accession number GSE142112).

## Acknowledgments

We would like to thank the Southall lab and Imperial College fly community for help and feedback on this project. We thank Mareike Jordan, Kristina Kleinschnitz and Nina Viessmann for generating and characterizing the Cap-G-EGFP and Barren-EGFP CRISPR knock-in lines. We are particularly grateful to Alicia Estacio-Gomez, Owen Marshall, and Seth Cheetham for critical reading of this manuscript. We would like to thank Andrea Brand for generously providing antibodies used in these experiments. For fly stocks, we thank the Bloomington *Drosophila* Stock Center (NIH P40OD018537). The Facility for Imaging by Light Microscopy (FILM) at Imperial College London is part-supported by funding from the Wellcome Trust (grant 104931/Z/14/Z) and BBSRC (grant BB/L015129/1). This work was funded by a Wellcome Trust Investigator grant 104567 to T.D.S, a BBSRC grant BB/P017924/1 to T.D.S. and G.N.A, and a BBSRC 1+3 DTP studentship (Amira Hassan).

## REFERENCES

1. Cajal, S., Sobre la aparición de las expansiones celulares en la médula embrionaria Gaceta Sanitaria de Barcelona 1890. 12: p. 413–419.

2. Southall, T.D., et al., Dedifferentiation of neurons precedes tumor formation in Lola mutants. Dev Cell, 2014. 28(6): p. 685–96.

3. Zacharioudaki, E., J. Falo Sanjuan, and S. Bray, Mi-2/NuRD complex protects stem cell progeny from mitogenic Notch signaling. Elife, 2019. 8.

4. Zeisel, A., et al., Molecular Architecture of the Mouse Nervous System. Cell, 2018. 174(4): p. 999–1014 e22.

5. Berger, C., et al., FACS Purification and Transcriptome Analysis of Drosophila Neural Stem Cells Reveals a Role for Klumpfuss in Self-Renewal. Cell Reports, 2012. 2(2): p. 407–418.

6. Zhang, S., et al., Open chromatin dynamics reveals stage-specific transcriptional networks in hiPSC-based neurodevelopmental model. Stem Cell Res, 2018. 29: p. 88–98.

7. Bonev, B., et al., Multiscale 3D Genome Rewiring during Mouse Neural Development. Cell, 2017. 171(3): p. 557–572 e24.

8. Chathoth, K.T. and N.R. Zabet, Chromatin architecture reorganization during neuronal cell differentiation in Drosophila genome. Genome research, 2019. 29(4): p. 613–625.

9. Aughey, G.N., et al., CATaDa reveals global remodelling of chromatin accessibility during stem cell differentiation in vivo. Elife, 2018. 7.

10. Yuen, K.C. and J.L. Gerton, Taking cohesin and condensin in context. PLoS Genet, 2018. 14(1): p. e1007118.

11. Hirano, T., Condensin-Based Chromosome Organization from Bacteria to Vertebrates. Cell, 2016. 164(5): p. 847–57.

12. Ganji, M., et al., Real-time imaging of DNA loop extrusion by condensin. Science, 2018.

13. Kschonsak, M., et al., Structural Basis for a Safety-Belt Mechanism That Anchors Condensin to Chromosomes. Cell, 2017. 171(3): p. 588–600 e24.

14. Terakawa, T., et al., The condensin complex is a mechanochemical motor that translocates along DNA. Science, 2017. 358(6363): p. 672–676.

15. Oliveira, R.A., P.A. Coelho, and C.E. Sunkel, The condensin I subunit Barren/CAP-H is essential for the structural integrity of centromeric heterochromatin during mitosis. Mol Cell Biol, 2005. 25(20): p. 8971–84.

16. Wallace, H.A. and G. Bosco, Condensins and 3D Organization of the Interphase Nucleus. Curr Genet Med Rep, 2013. 1(4): p. 219–229.

17. Dowen, J.M. and R.A. Young, SMC complexes link gene expression and genome architecture. Current opinion in genetics & development, 2014. 25: p. 131–137.

18. Nakazawa, N., et al., RNA pol II transcript abundance controls condensin accumulation at mitotically up-regulated and heat-shock-inducible genes in fission yeast. Genes Cells, 2015. 20(6): p. 481–99.

19. Dowen, J.M., et al., Multiple structural maintenance of chromosome complexes at transcriptional regulatory elements. Stem Cell Reports, 2013. 1(5): p. 371–8.

20. D’Ambrosio, C., et al., Identification of cis-acting sites for condensin loading onto budding yeast chromosomes. Genes Dev, 2008. 22(16): p. 2215–27.

21. Kranz, A.L., et al., Genome-wide analysis of condensin binding in Caenorhabditis elegans. Genome Biol, 2013. 14(10): p. R112.

22. Kim, J.H., et al., Condensin I associates with structural and gene regulatory regions in vertebrate chromosomes. Nat Commun, 2013. 4: p. 2537.

23. Yuen, K.C., B.D. Slaughter, and J.L. Gerton, Condensin II is anchored by TFIIIC and H3K4me3 in the mammalian genome and supports the expression of active dense gene clusters. Science advances, 2017. 3(6): p. e1700191.

24. Li, W., et al., Condensin I and II complexes license full estrogen receptor α-dependent enhancer activation. Molecular cell, 2015. 59(2): p. 188–202.

25. Li, L., et al., Widespread rearrangement of 3D chromatin organization underlies polycomb-mediated stress-induced silencing. Mol Cell, 2015. 58(2): p. 216–31.

26. Swygert, S.G., et al., Condensin-Dependent Chromatin Compaction Represses Transcription Globally during Quiescence. Mol Cell, 2019. 73(3): p. 533–546 e4.

27. Rawlings, J.S., et al., Chromatin condensation via the condensin II complex is required for peripheral T-cell quiescence. The EMBO journal, 2011. 30(2): p. 263–276.

28. Dej, K.J., C. Ahn, and T.L. Orr-Weaver, Mutations in the Drosophila condensin subunit dCAP-G: defining the role of condensin for chromosome condensation in mitosis and gene expression in interphase. Genetics, 2004. 168(2): p. 895–906.

29. Savvidou, E., et al., Drosophila CAP-D2 is required for condensin complex stability and resolution of sister chromatids. J Cell Sci, 2005. 118(Pt 11): p. 2529–43.

30. Cobbe, N., E. Savvidou, and M.M. Heck, Diverse mitotic and interphase functions of condensins in Drosophila. Genetics, 2006. 172(2): p. 991–1008.

31. Lupo, R., et al., Drosophila chromosome condensation proteins Topoisomerase II and Barren colocalize with Polycomb and maintain Fab-7 PRE silencing. Molecular cell, 2001. 7(1): p. 127–136.

32. Hartl, T.A., H.F. Smith, and G. Bosco, Chromosome alignment and transvection are antagonized by condensin II. Science, 2008. 322(5906): p. 1384–1387.

33. Meyer, B.J., Targeting X chromosomes for repression. Current opinion in genetics & development, 2010. 20(2): p. 179–189.

34. Xu, Y., et al., MTB, the murine homolog of condensin II subunit CAP-G2, represses transcription and promotes erythroid cell differentiation. Leukemia, 2006. 20(7): p. 1261–9.

35. Paul, M.R., et al., Condensin Depletion Causes Genome Decompaction Without Altering the Level of Global Gene Expression in Saccharomyces cerevisiae. Genetics, 2018. 210(1): p. 331–344.

36. Nishide, K. and T. Hirano, Overlapping and non-overlapping functions of condensins I and II in neural stem cell divisions. PLoS genetics, 2014. 10(12): p. e1004847.

37. Woodward, J., et al., Condensin II mutation causes T-cell lymphoma through tissue-specific genome instability. Genes & development, 2016. 30(19): p. 2173–2186.

38. Hocquet, C., et al., Condensin controls cellular RNA levels through the accurate segregation of chromosomes instead of directly regulating transcription. Elife, 2018. 7.

39. Abdennur, N., et al., Condensin II inactivation in interphase does not affect chromatin folding or gene expression. BioRxiv, 2018: p. 437459.

40. Longworth, M.S., et al., A shared role for RBF1 and dCAP-D3 in the regulation of transcription with consequences for innate immunity. PLoS Genet, 2012. 8(4): p. e1002618.

41. Herzog, S., et al., Functional dissection of the Drosophila melanogaster condensin subunit Cap-G reveals its exclusive association with condensin I. PLoS Genet, 2013. 9(4): p. e1003463.

42. Leader, D.P., et al., FlyAtlas 2: a new version of the Drosophila melanogaster expression atlas with RNA-Seq, miRNA-Seq and sex-specific data. Nucleic acids research, 2017. 46(D1): p. D809–D815.

43. Oliveira, R.A., S. Heidmann, and C.E. Sunkel, Condensin I binds chromatin early in prophase and displays a highly dynamic association with Drosophila mitotic chromosomes. Chromosoma, 2007. 116(3): p. 259–74.

44. Caussinus, E., O. Kanca, and M. Affolter, Fluorescent fusion protein knockout mediated by anti-GFP nanobody. Nat Struct Mol Biol, 2011. 19(1): p. 117–21.

45. Jager, H., M. Rauch, and S. Heidmann, The Drosophila melanogaster condensin subunit Cap-G interacts with the centromere-specific histone H3 variant CID. Chromosoma, 2005. 113(7): p. 350–61.

46. Berger, C., et al., The commonly used marker ELAV is transiently expressed in neuroblasts and glial cells in the Drosophila embryonic CNS. Developmental dynamics: an official publication of the American Association of Anatomists, 2007. 236(12): p. 3562–3568.

47. Giet, R. and D.M. Glover, Drosophila aurora B kinase is required for histone H3 phosphorylation and condensin recruitment during chromosome condensation and to organize the central spindle during cytokinesis. J Cell Biol, 2001. 152(4): p. 669–82.

48. Seipold, S., et al., Non-SMC condensin I complex proteins control chromosome segregation and survival of proliferating cells in the zebrafish neural retina. BMC Dev Biol, 2009. 9: p. 40.

49. Lanson, N.A., Jr., et al., A Drosophila model of FUS-related neurodegeneration reveals genetic interaction between FUS and TDP-43. Hum Mol Genet, 2011. 20(13): p. 2510–23.

50. Gargano, J.W., et al., Rapid iterative negative geotaxis (RING): a new method for assessing age-related locomotor decline in Drosophila. Exp Gerontol, 2005. 40(5): p. 386–95.

51. Lake, C.M., et al., The development of a monoclonal antibody recognizing the Drosophila melanogaster phosphorylated histone H2A variant (gamma-H2AV). G3 (Bethesda), 2013. 3(9): p. 1539–43.

52. Southall, T.D., et al., Cell-type-specific profiling of gene expression and chromatin binding without cell isolation: assaying RNA Pol II occupancy in neural stem cells. Dev Cell, 2013. 26(1): p. 101–12.

53. Aughey, G.N., S.W. Cheetham, and T.D. Southall, DamID as a versatile tool for understanding gene regulation. Development, 2019. 146(6): p. dev173666.

54. Chia, W., W.G. Somers, and H. Wang, Drosophila neuroblast asymmetric divisions: cell cycle regulators, asymmetric protein localization, and tumorigenesis. J Cell Biol, 2008. 180(2): p. 267–72.

55. Homem, C.C. and J.A. Knoblich, Drosophila neuroblasts: a model for stem cell biology. Development, 2012. 139(23): p. 4297–310.

56. Rivera, J., et al., REDfly: the transcriptional regulatory element database for Drosophila. Nucleic acids research, 2018. 47(D1): p. D828–D834.

57. Filion, G.J., et al., Systematic protein location mapping reveals five principal chromatin types in Drosophila cells. Cell, 2010. 143(2): p. 212–224.

58. Marshall, O.J. and A.H. Brand, Chromatin state changes during neural development revealed by in vivo cell-type specific profiling. Nat Commun, 2017. 8(1): p. 2271.

59. Gallegos, D.A., et al., Chromatin Regulation of Neuronal Maturation and Plasticity. Trends Neurosci, 2018. 41(5): p. 311–324.

60. Venkatesh, I., et al., Epigenetic profiling reveals a developmental decrease in promoter accessibility during cortical maturation in vivo. Neuroepigenetics, 2016. 8: p. 19–26.

61. Wang, J.T., et al., Disease gene candidates revealed by expression profiling of retinal ganglion cell development. J Neurosci, 2007. 27(32): p. 8593–603.

62. Froldi, F., et al., The transcription factor Nerfin-1 prevents reversion of neurons into neural stem cells. Genes Dev, 2015. 29(2): p. 129–43.

63. Palmisano, I. and S. Di Giovanni, Advances and Limitations of Current Epigenetic Studies Investigating Mammalian Axonal Regeneration. Neurotherapeutics, 2018. 15(3): p. 529–540.

64. Brugger, V., et al., Delaying histone deacetylase response to injury accelerates conversion into repair Schwann cells and nerve regeneration. Nat Commun, 2017. 8: p. 14272.

65. Lau, A.C. and G. Csankovszki, Condensin-mediated chromosome organization and gene regulation. Front Genet, 2014. 5: p. 473.

66. Aono, N., et al., Cnd2 has dual roles in mitotic condensation and interphase. Nature, 2002. 417(6885): p. 197–202.

67. Chen, E.S., T. Sutani, and M. Yanagida, Cti1/C1D interacts with condensin SMC hinge and supports the DNA repair function of condensin. Proc Natl Acad Sci U S A, 2004. 101(21): p. 8078–83.

68. Heale, J.T., et al., Condensin I interacts with the PARP-1-XRCC1 complex and functions in DNA single-strand break repair. Mol Cell, 2006. 21(6): p. 837–48.

69. Van Bortle, K., et al., Insulator function and topological domain border strength scale with architectural protein occupancy. Genome biology, 2014. 15(5): p. R82.

70. McDonel, P., et al., Clustered DNA motifs mark X chromosomes for repression by a dosage compensation complex. Nature, 2006. 444(7119): p. 614.

71. Paul, M.R., A. Hochwagen, and S. Ercan, Condensin action and compaction. Curr Genet, 2019. 65(2): p. 407–415.

72. Pauli, A., et al., Cell-type-specific TEV protease cleavage reveals cohesin functions in Drosophila neurons. Dev Cell, 2008. 14(2): p. 239–51.

73. Albertson, R., et al., Scribble protein domain mapping reveals a multistep localization mechanism and domains necessary for establishing cortical polarity. J Cell Sci, 2004. 117(Pt 25): p. 6061–70.

74. Yang, L., et al., Single molecule fluorescence in situ hybridisation for quantitating post-transcriptional regulation in Drosophila brains. Methods, 2017. 126: p. 166–176.

75. Schindelin, J., et al., Fiji: an open-source platform for biological-image analysis. Nature methods, 2012. 9(7): p. 676.

76. De Chaumont, F., et al., Icy: an open bioimage informatics platform for extended reproducible research. Nature methods, 2012. 9(7): p. 690.

77. Olivo-Marin, J.-C., Extraction of spots in biological images using multiscale products. Pattern recognition, 2002. 35(9): p. 1989–1996.

78. Berg, S., et al., ilastik: interactive machine learning for (bio) image analysis. Nature Methods, 2019: p. 1–7.

79. Nichols, C.D., J. Becnel, and U.B. Pandey, Methods to assay Drosophila behavior. JoVE (Journal of Visualized Experiments), 2012(61): p. e3795.

80. Bischof, J., et al., An optimized transgenesis system for Drosophila using germ-line-specific phiC31 integrases. Proc Natl Acad Sci U S A, 2007. 104(9): p. 3312–7.

81. Gibson, D.G., et al., Chemical synthesis of the mouse mitochondrial genome. Nat Methods, 2010. 7(11): p. 901–3.

82. Marshall, O.J., et al., Cell-type-specific profiling of protein–DNA interactions without cell isolation using targeted DamID with next-generation sequencing. Nature protocols, 2016. 11(9): p. 1586.

83. Heinz, S., et al., Simple combinations of lineage-determining transcription factors prime cis-regulatory elements required for macrophage and B cell identities. Molecular cell, 2010. 38(4): p. 576–589.

84. Scientific, T.F. 2016; Available from: https://tools.thermofisher.com/content/sfs/manuals/trizol_reagent.pdf.

85. Dobin, A., et al., STAR: ultrafast universal RNA-seq aligner. Bioinformatics, 2013. 29(1): p. 15–21.

86. Liao, Y., G.K. Smyth, and W. Shi, The R package Rsubread is easier, faster, cheaper and better for alignment and quantification of RNA sequencing reads. Nucleic acids research, 2019. 47(8): p. e47–e47.

87. Love, M.I., W. Huber, and S. Anders, Moderated estimation of fold change and dispersion for RNA-seq data with DESeq2. Genome biology, 2014. 15(12): p. 550.

88. Blighe, K., EnhancedVolcano: Publication-ready volcano plots with enhanced colouring and labeling. R package version 1.2. 0. 2019.

89. McKinney, W. Data structures for statistical computing in python. in Proceedings of the 9th Python in Science Conference. 2010. Austin, TX.

90. Quinlan, A.R. and I.M. Hall, BEDTools: a flexible suite of utilities for comparing genomic features. Bioinformatics, 2010. 26(6): p. 841–2.

91. Ramírez, F., et al., deepTools2: a next generation web server for deep-sequencing data analysis. Nucleic acids research, 2016. 44(W1): p. W160–W165.

92. Yu, G., et al., clusterProfiler: an R package for comparing biological themes among gene clusters. Omics: a journal of integrative biology, 2012. 16(5): p. 284–287.

