## Supplemental figures for "Condensin I subunit Cap-G is essential for proper gene expression during the maturation of post-mitotic neurons"

### **Supplementary information**

#### **Contents:**

**Figure 1 – figure supplement 1.** *Cap-G<sup>EGFP</sup>* expression in neurons.

**Figure 2 – figure supplement 1.** Characterisation of Cap-G knockdown with *elav-GAL4*.

**Figure 2 – figure supplement 2.** Apoptosis in *elav-KD*.

**Figure 4 – figure supplement 1.** Analysis of Cap-G binding.

**Figure 5 – figure supplement 1.** Analysis of RNA-seq data.

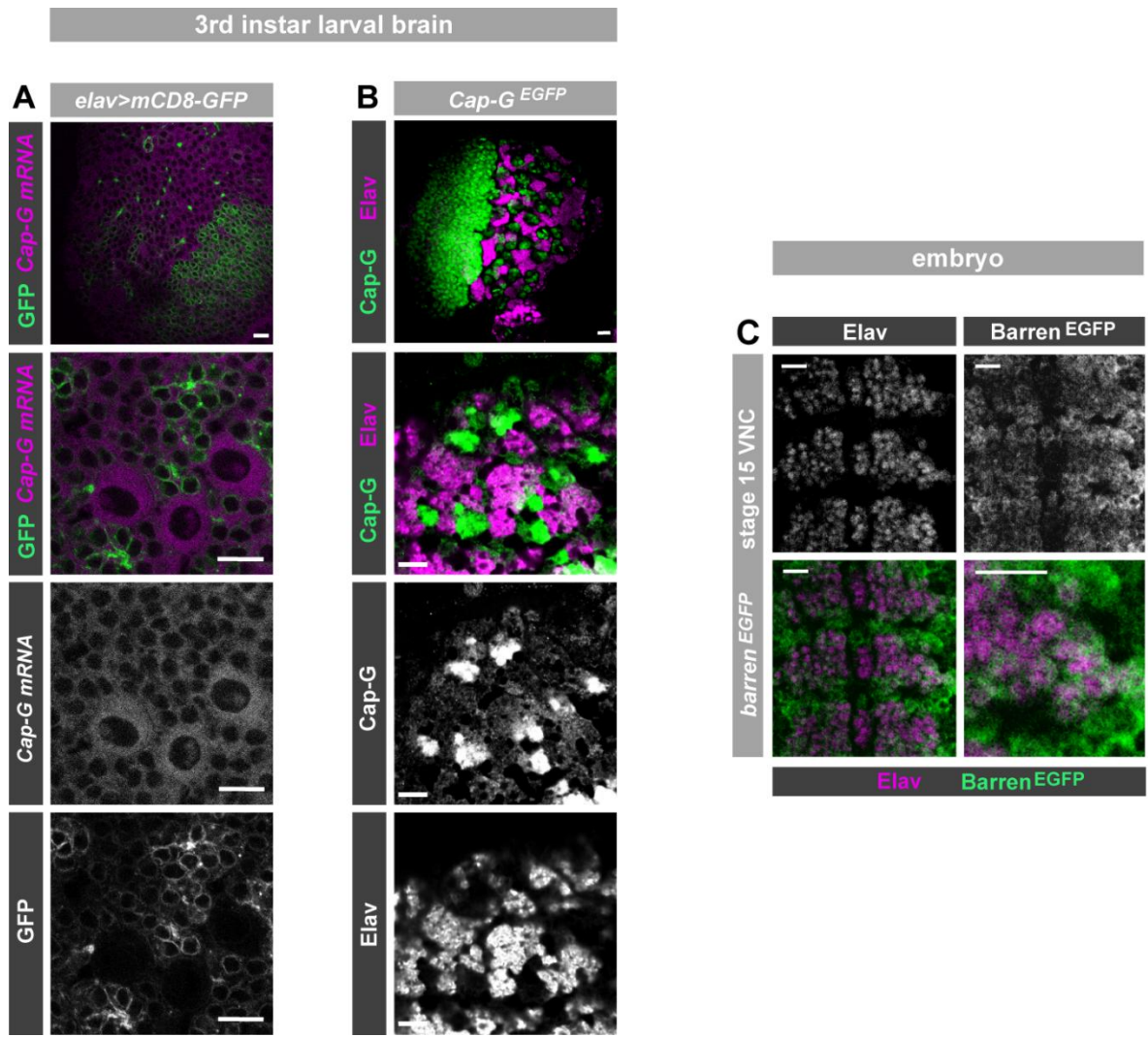

**Figure 1 – figure supplement 1. *Cap-G<sup>EGFP</sup>* expression in neurons.** **A)** Optic lobe of 3<sup>rd</sup> instar larvae. *Cap-G mRNA* is ubiquitously expressed in the lobes. mCD8-GFP marks neuronal membrane. Zoom in of central brain section shows expression of *Cap-G mRNA* in neurons marked by elav-driven GFP expression. **B)** Optic lobe of 3<sup>rd</sup> instar *Cap-G<sup>EGFP</sup>* larvae. Cap-G is strongly present in the optic proliferation centre and the central brain. Zoom in on CB shows Cap-G is present in neuronal nuclei as it overlaps with marker Elav. **C)** *Barren<sup>EGFP</sup>* embryo (stage 15). Barren<sup>EGFP</sup> is present in all cells of the VNC, including Elav positive neurons. All images show ventral view of embryonic VNC (anterior -top). Scale bars = 10 μm.

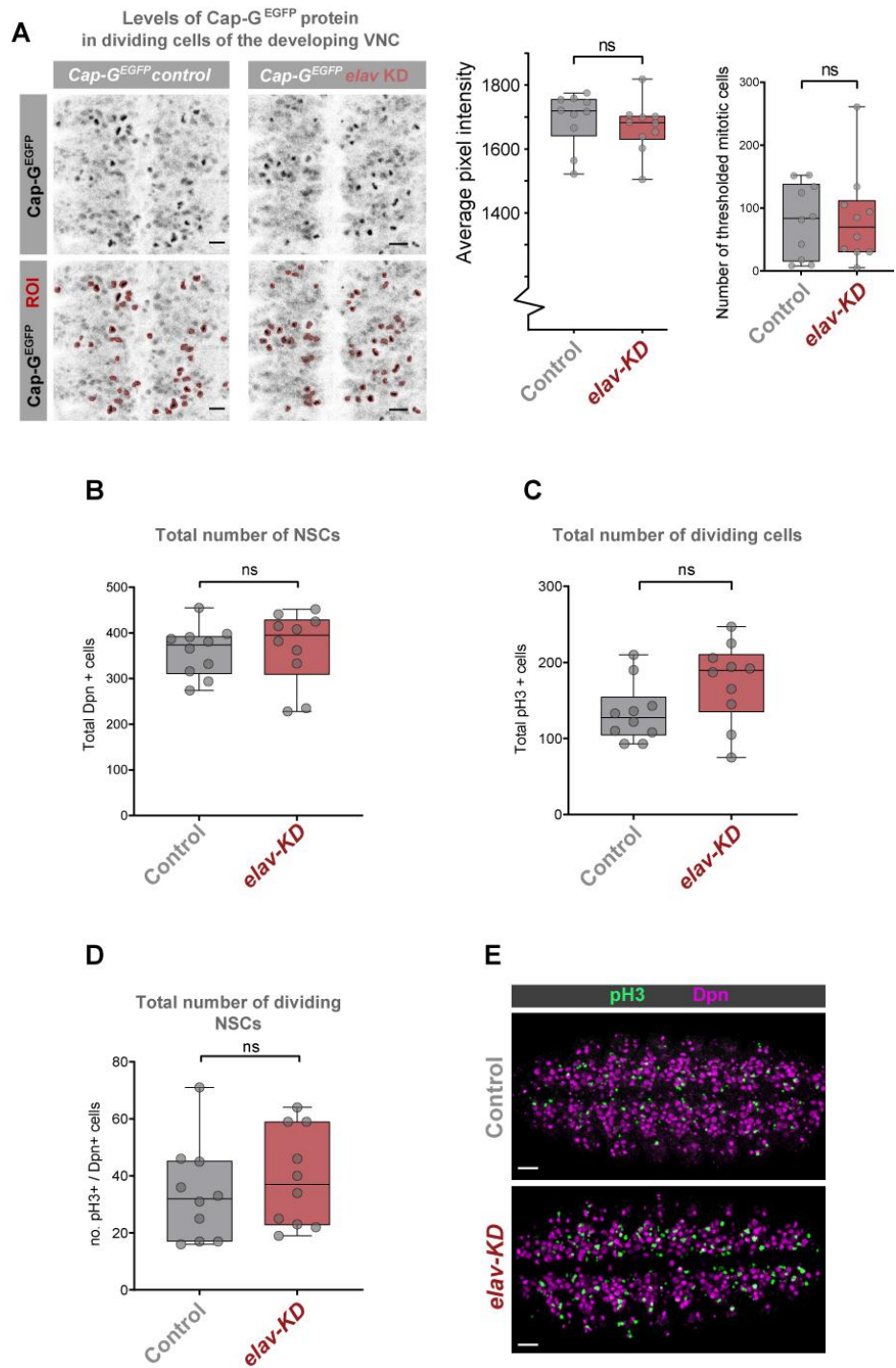

**Figure 2 - figure supplement 1. Characterisation of Cap-G knockdown with *elav-GAL4* (*elav-KD*).** **A)** Quantification of Cap-G<sup>EGFP</sup> protein levels in dividing cells in *Cap-G<sup>EGFP</sup>* control and *elav-KD* embryonic VNC (stage 14 /15- anterior top). Pixel intensity thresholding was performed to produce images in which dividing cells were defined as regions of interest (ROI) whilst neurons (lower pixel intensity) were excluded from the analysis. The average pixel intensity of dividing cells is calculated for >30 cells/ embryo, with 10 biological replicates per genotype. No significant difference in Cap-G<sup>EGFP</sup> levels, t-test ($p > 0.5$). Number of mitotic cells (ROIs) remains constant. **B-D)** Quantification of NSCs numbers and cell division in *Cap-G<sup>EGFP</sup>* control and *elav-KD* individuals. Z-stacks of embryonic VNC stage 15 for *Cap-G<sup>EGFP</sup>* and *elav-KD* were used. **B)** Quantification of total number of NSCs marked by Dpn. **C)** Quantification of total number of mitotic cells marked by pH3. **D)** Quantification of total number of dividing NSCs as number of pH3+/Dpn+ cells. 10 replicates per genotype. No significant difference as per t-test ($p > 0.05$). **E)** Maximum Z-projection of stage 15 embryonic VNC for both *Cap-G<sup>EGFP</sup>* and *elav-KD* showing quantified NSCs (Dpn) and dividing cells (pH3). Scale bar 10 μm.

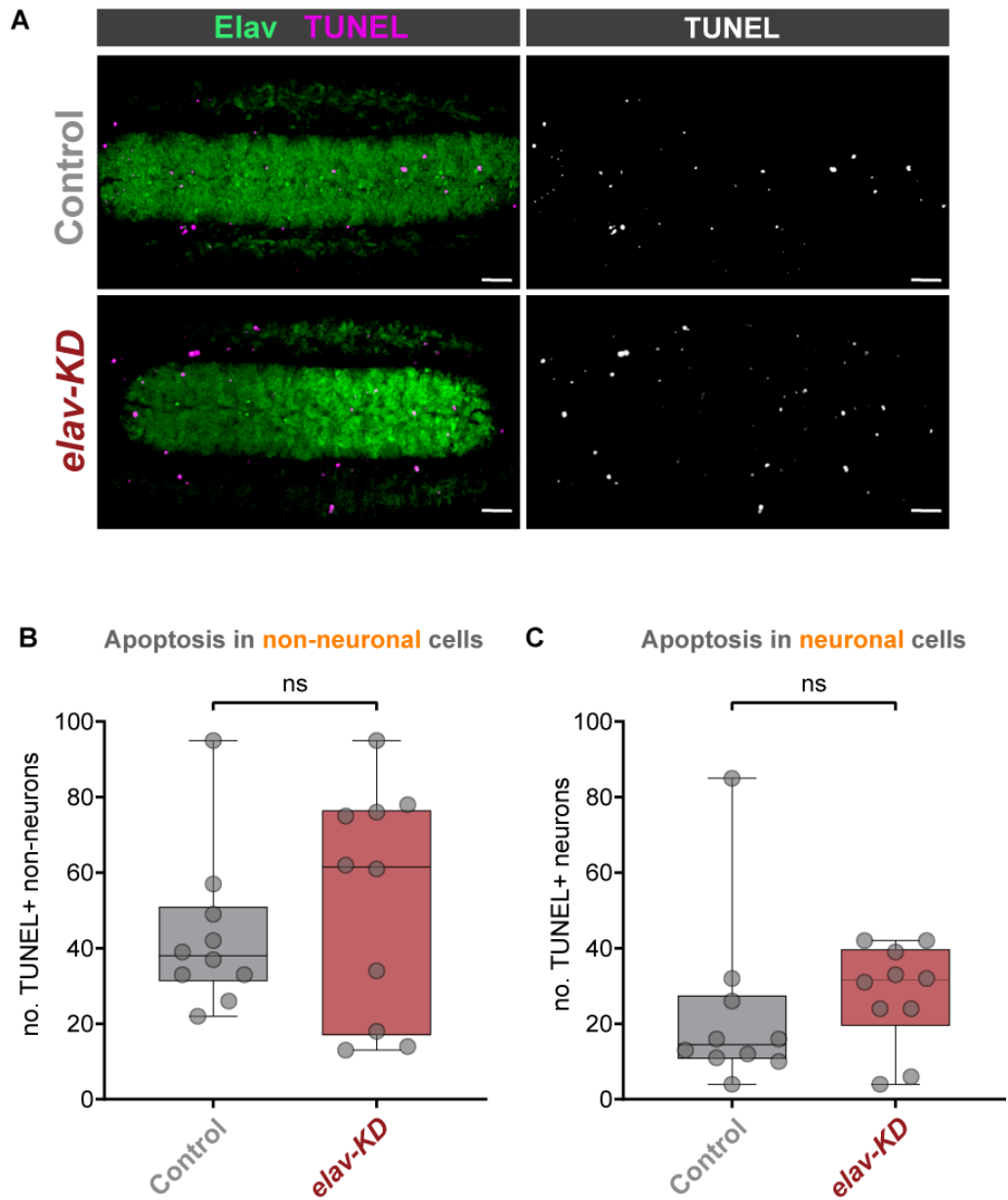

**Figure 2 – figure supplement 2. Apoptosis in *elav-KD*.** **A)** Staining for apoptotic cells in embryonic VNC stage 16 for *Cap-G<sup>EGFP</sup>* (control) and *elav-KD*. Neurons marked in green by Elav and apoptotic cells in magenta by TUNEL staining. Anterior left, posterior right. Maximum Z-projection. Scale bar 10 μm. **B-C)** Quantification of apoptosis in embryonic VNC stage 15/16 for *Cap-G<sup>EGFP</sup>* and *elav-KD*. 10 replicates per genotype. **B)** No significant difference in number of apoptotic non-neuronal cells (TUNEL +, Elav -) between *Cap-G<sup>EGFP</sup>* control and *elav-KD*. **C)** No significant difference in apoptosis in neurons (TUNEL+, Elav +). T-test $p > 0.05$.

**A**

**Correlation between accessible chromatin (AC) and Cap-G binding**

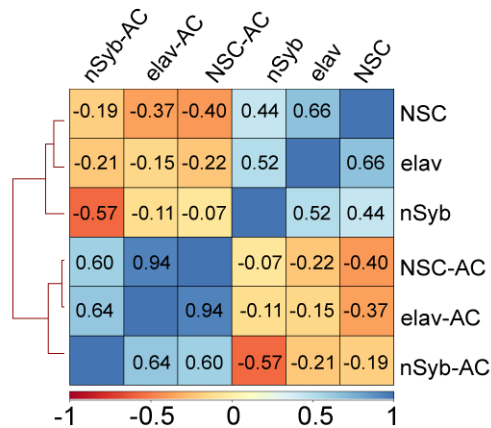

**B**

**Principal Component Analysis between AC and Cap-G binding**

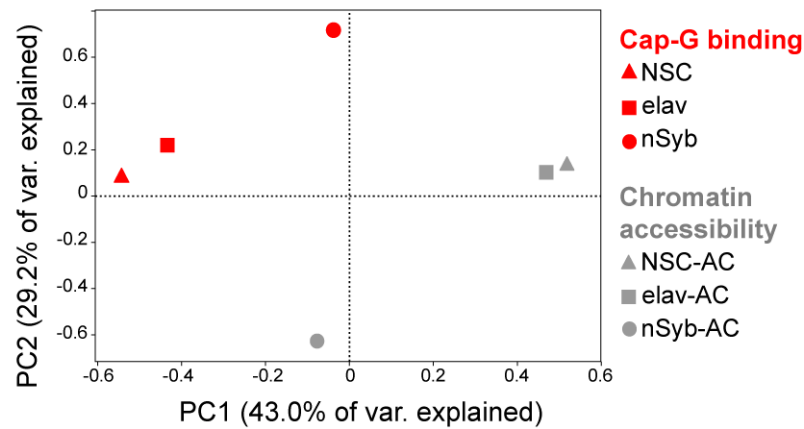

**Figure 4 – figure supplement 1. Analysis of Cap-G binding.** **A)** Correlation heatmap between Cap-G binding and Accessible Chromatin (AC) per cell type. Cap-G binding negatively correlates with AC in each cell type. Numbers represent Spearman's rank correlation coefficient. Cap-G binding and AC samples cluster separately by k-means. **B)** PCA of Cap-G binding and AC per cell type. Cap-G binding (red) and AC (grey) cluster separately. Elav and NSC cluster in proximity of each other reflecting their greater similarity, whilst nSyb is more distant, observed in both AC and Cap-G binding samples.

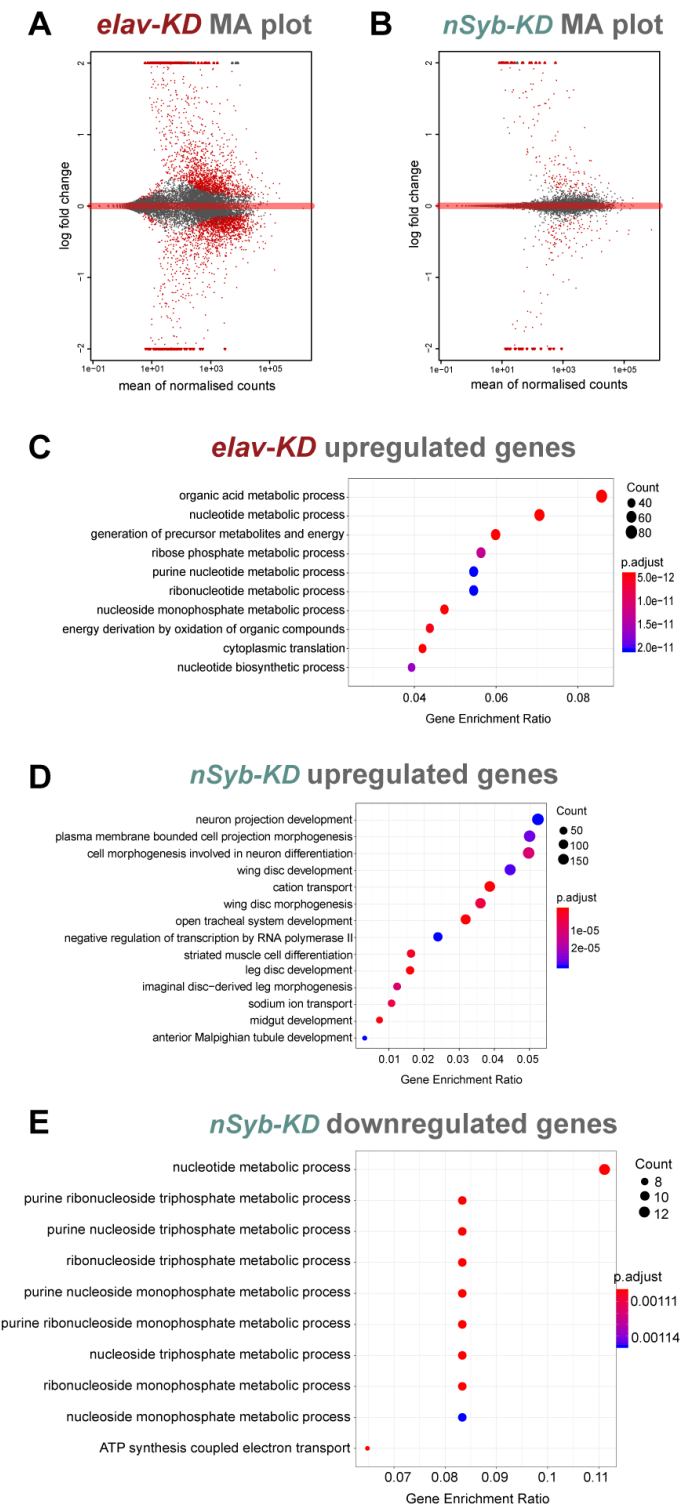

**Figure 5 – figure supplement 1. Analysis of RNA-seq data. A-B)** MA plot of RNAseq datasets for *elav-KD* and *nSyb-KD* respectively showing mean of normalised counts and log<sub>2</sub> fold change values, significant genes marked in red. **C-E)** Enriched GO terms analysis for differentially expressed genes, gene enrichment ratio shown, circle size correlates with number of genes detected and colour with p-adjust value. **C)** Upregulated genes in *elav-KD* showing generic, non-CNS specific terms GO terms. **D)** Upregulated genes in *nSyb-KD* showing generic, non-CNS specific terms. **E)** Downregulated genes in *nSyb-KD* showing metabolic-related GO terms.
